# Uncovering Interpretable Fine-Grained Phenotypes of Subcellular Dynamics through Unsupervised Self-Training of Deep Neural Networks

**DOI:** 10.1101/2021.05.25.445699

**Authors:** Chuangqi Wang, Hee June Choi, Lucy Woodbury, Kwonmoo Lee

## Abstract

Live cell imaging provides unparallel insights into dynamic cellular processes across spatiotemporal scales. Despite its potential, the inherent spatiotemporal heterogeneity within live cell imaging data often obscures critical mechanical details underlying cellular dynamics. Uncovering fine-grained phenotypes of live cell dynamics is pivotal for precise understandings of the heterogeneity of physiological and pathological processes. However, this endeavor introduces formidable technical challenges to unsupervised machine learning, demanding the extraction of features that can faithfully preserve heterogeneity, effectively discriminate between different molecularly perturbed states, and provide interpretability. While deep learning shows promise in extracting useful features from large datasets, it often falls short in producing such high-fidelity features, especially in unsupervised learning settings. To tackle these challenges, we present DeepHACX (Deep phenotyping of Heterogeneous Activities of Cellular dynamics with eXplanations), a self-training deep learning framework designed for fine-grained and interpretable phenotyping. This framework seamlessly integrates an unsupervised teacher model with interpretable features to facilitate feature learning in a student deep neural network (DNN). Significantly, it incorporates an autoencoder-based regularizer, termed SENSER (SENSitivity-enhancing autoEncoding Regularizer), designed to prompt the student DNN to maximize the heterogeneity associated with molecular perturbations. This approach enables the acquisition of features that not only discriminate between different molecularly perturbed states but also faithfully preserve the heterogeneity linked to these perturbations. In our study, DeepHACX successfully delineated fine-grained phenotypes within the heterogeneous protrusion dynamics of migrating epithelial cells, uncovering specific responses to pharmacological perturbations. Remarkably, DeepHACX adeptly captured a minimal number of highly interpretable features uniquely linked to these fine-grained phenotypes, each corresponding to specific temporal intervals crucial for their manifestation. This unique capability positions DeepHACX as a valuable tool for investigating diverse cellular dynamics and comprehensively studying their heterogeneity.

## Introduction

Live cell imaging offers unique insights into dynamic cellular processes across various spatiotemporal scales^1–4^. Recent advancements in microscopy technology have enabled the acquisition of extensive live cell image data at high resolutions. However, conventional ensemble-averaging approaches may overlook critical mechanistic details due to spatiotemporal heterogeneity in live cell data^4–6^. This becomes particularly problematic when specific molecular perturbations induce responses in only a limited subcellular region of a rare cellular subpopulation, a phenomenon often overlooked in ensemble averaging without the deconvolution of heterogeneity.

Although computational image analysis and machine learning have been extensively utilized for cellular heterogeneity analysis, the majority of these efforts have concentrated on static images^7–9^. The investigation into phenotypic heterogeneity in live cell images is still in its nascent stages^4,6,10–13^,necessitating the development of novel features that accurately represent subcellular and cellular temporal dynamics for effective deconvolution of heterogeneous phenotypes. Specifically, the exploration of live cell fine-grained phenotypes, referred to as deep phenotyping for brevity in this work^14,15^, has been hampered by significant technical challenges. This endeavor requires the detection of nuanced differences in cellular dynamics with limited prior knowledge.

In deep phenotyping, handling high-dimensional datasets requires informative features that effectively reduce dimensionality while retaining crucial information, addressing challenges associated with the curse of dimensionality^16,17^. Traditional feature engineering involves domain-specific hand-crafted designs, which may not be directly applicable to deep phenotyping. Deep learning has emerged as a promising alternative due to its exceptional ability to learn comprehensive representative features directly from large raw datasets^18,19^. However, conventional methods for feature learning encounter challenges when addressing the complexities of deep phenotyping. Conventional neural network training, focusing on improving classification accuracy, often produces features that might not comprehensively capture the entire spectrum of intra-class heterogeneity, particularly when dealing with limited data prone to overfitting. Conversely, autoencoders^20^ can identify extensive features across samples, but these features may lack the discriminatory power required for recognizing specific cellular or disease states^21,22^. Additionally, they might include considerable irrelevant heterogeneity, compromising their effectiveness in unsupervised learning. Moreover, an ongoing challenge lies in making the learned features interpretable to domain experts^23,24^. Interpreting these features within the context of cellular dynamics remains an active area of research, as it enables a deeper understanding of the underlying mechanisms^25^. These limitations underscore the need for more advanced approaches in feature learning capable of preserving relevant heterogeneity and ensuring accurate discrimination while enhancing their interpretability.

To overcome these challenges, we developed DeepHACX (Deep phenotyping of Heterogeneous Activities of Cellular dynamics with eXplanations), an unsupervised self-training deep learning framework specifically designed for identifying interpretable fine-grained phenotypes of live cell dynamics. DeepHACX leverages the power of DNNs to unravel the phenotypic heterogeneity present in subcellular time-series datasets. In this self-training framework, we employ an unsupervised teacher model to guide the learning process of a student DNN. The student DNN is tailored to capture features that faithfully preserve the heterogeneity stemming from the effects of molecular perturbations. Our approach ensures both relevant heterogeneity and interpretability of the learned features, enabling a deeper comprehension of the underlying cellular dynamics.

In this study, we employed DeepHACX to analyze the dynamic behavior of migrating epithelial cells’ leading edges, which undergo protrusion and retraction cycles. Cell protrusion is crucial for tissue regeneration^26,27^, cancer invasiveness and metastasis^28–30^, and microenvironmental surveillance of leukocytes^31^. Coordinated protrusion and retraction are linked to the metastatic potential of cancer cells^32^ and phenotypic switching of cell migration^33^, making the analysis of morphodynamic biomarkers relevant for diagnosing breast and prostate cancer^34^. However, dissecting these dynamics has been challenging due to the significant heterogeneity^5,6,35^. DeepHACX successfully identified fine-grained phenotypes from the highly heterogeneous and non-stationary edge dynamics of migrating epithelial cells. It revealed specific responses to pharmacological perturbations in the form of “bursting” and “accelerating” protrusions. Intriguingly, DeepHACX generated distinct, minimal features associated with these nuanced phenotypes, enhancing interpretability by precisely identifying the critical temporal intervals governing their manifestation. Furthermore, we demonstrated the ability to define single-cell phenotypes based on these protrusions. This remarkable capability solidifies DeepHACX as a valuable tool for exploring diverse cellular dynamics and understanding their heterogeneous responses to molecular perturbations.

## Results

### DeepHACX: an interpretable self-training framework for fine-grained phenotyping of subcellular dynamics

DNNs are adept at learning comprehensive features from raw datasets, making them well-suited for the discovery of phenotypes. However, the multitude of features generated by DNNs often lack interpretability and the essential heterogeneity required for their effective application in fine-grained phenotyping. To address this limitation, we propose a machine learning framework, DeepHACX (**Fig. 1a**), which integrates unsupervised learning with self-training. This allows a student DNN to acquire a concise and interpretable set of informative features capturing the temporal dynamics, guided by a teacher model. Self-training, a widely recognized semi-supervised learning method, involves a teacher model trained on a restricted labeled dataset providing pseudo-labels for unlabeled data to train a student model^36–39^. While this approach has proven effective in supervised learning with limited labeled and abundant unlabeled datasets, we employ it here in an unsupervised context. Our teacher model is an unsupervised model utilizing straightforward interpretable features. Then, we introduce a novel regularization strategy for training the student DNN, aiming to extract informative features that preserve heterogeneity while discriminating molecularly perturbed states. This integration of interpretability and feature learning ensures the extraction of a minimal number of interpretable features, facilitating the identification of fine-grained phenotypes with distinct sensitivities to molecular perturbations.

**Figure 1.**
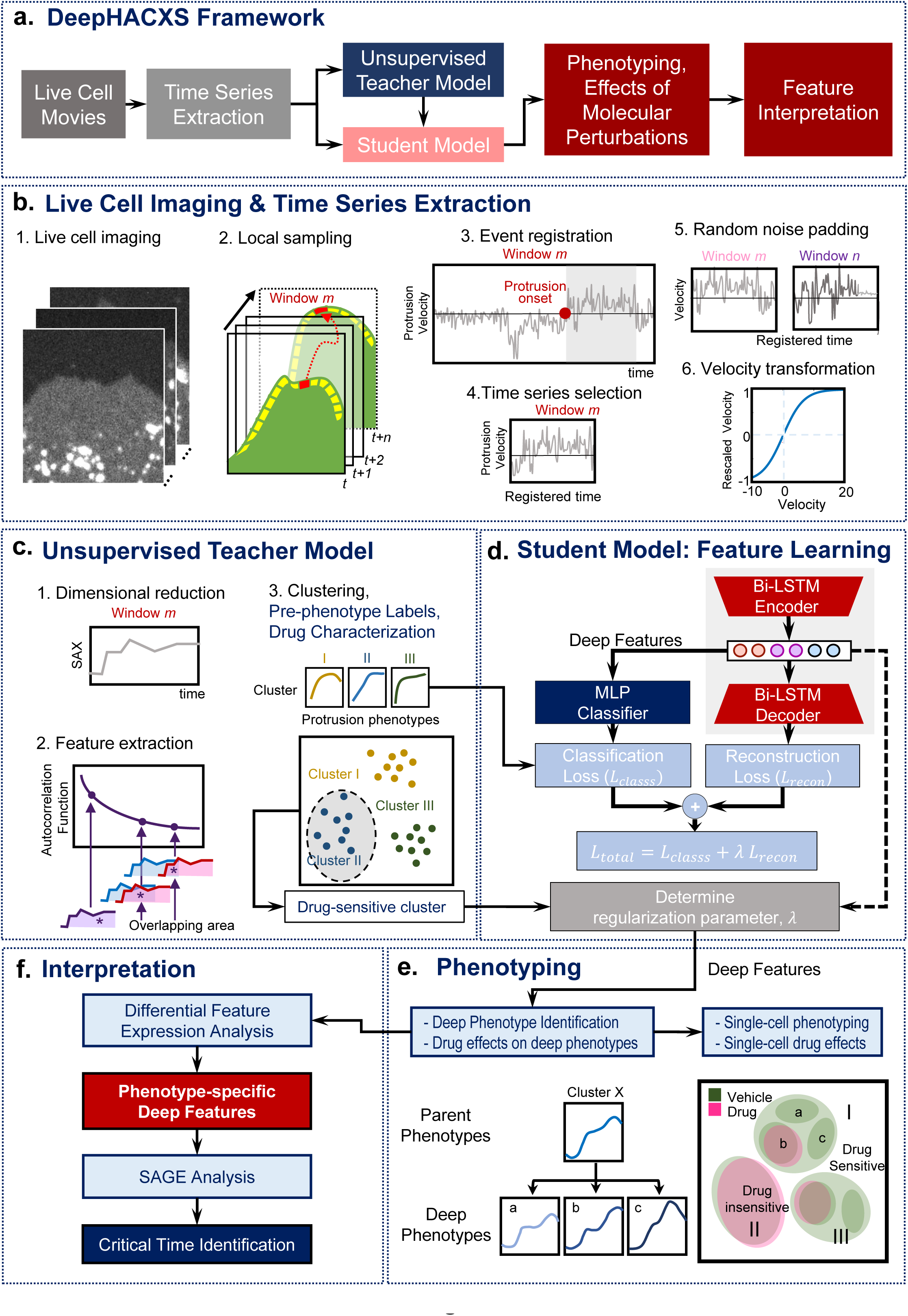
Schematic representation of the computational pipeline. (**a**) Self-training framework of DeepHACX. (**b**) Time series data generation and preprocessing. (1-2) Live cell imaging and local sampling. Time-lapse movies of the leading edge of migrating cells treated with/without different drugs were taken at 5 sec per frame, and then probing windows (500 x 500nm) were generated to track the cell edge movement and local velocities were calculated per window. (3-4). Event registration and time series selection. The velocity profile per window was divided into protrusion and retraction events based on protrusion onset. Then all the protrusion intervals with a length longer than 50 sec are selected, and the samples with longer protrusion duration are truncated to 250s for further analysis. (5-6) Random noise padding and velocity transformation. The raw velocity profiles were non-linearly scaled to [-1, 1] for the purpose of convenient training and eliminating the effect of larger values. (**c**) Pseudo-labels generated by the teacher model using interpretable features such as ACF. ACF-based clustering pipeline was applied to the collected data to identify the protrusion phenotypes and characterize their sensitivities of molecular perturbations. (**d**) Feature Learning by the student deep model integrating an BiLSTM-autoencoder and multiple-layer perceptron (MLP) classifier. The regularization parameter to balance the contributions of two branches was chosen to maximize the sensitivity of given molecular perturbations. The deep features from the bi-LSTM encoder were extracted for further phenotyping. (**e**) Phenotyping. Subcellular phenotypes were identified by clustering analysis using deep features. The identified phenotypes can be divided into deep phenotypes for more precise drug characterization. Subcellular phenotypes are used for single-cell phenotyping. (**f**) Feature Interpretation. We performed differential feature expression analysis to identify the deep features significantly associated with deep phenotypes. The SAGE analysis is applied to identify the time points critically linked to the significant deep features.

DeepHACX is a comprehensive framework comprising five key components: (i) an unsupervised teacher model using a predefined interpretable feature such as Autocorrelation Function (ACF), (ii) a student model specialized in deep feature learning, leveraging pseudo-labels generated by the teacher model, while being regularized to maximize the heterogeneity associated with molecular perturbations, (iii) fine-grained clustering and molecular perturbation analysis, (iv) analysis of feature interpretability, and (iv) single-cell phenotyping based on subcellular phenotypes. Our approach incorporates ACF-based clustering as the teacher model to derive pseudo-labels, seamlessly integrated into the student DNN for the automatic acquisition of interpretable features in deep phenotyping of live cells. Through the synergy of these components, DeepHACX offers a comprehensive and interpretable framework for uncovering fine-grained phenotypes and facilitating an in-depth understanding of complex cellular dynamics.

To prepare our sample videos, we imaged migrating PtK1 epithelial cells in control and drug-perturbed conditions (**Fig. 1b.1**). We then segmented the cell membrane boundary in each frame of each movie and divided the cell edges into small probing windows with a size of 500 by 500nm^5,40^. We acquired time series of protrusion velocities by averaging the velocities of pixels in each probing window (**Fig. 1b.2**). Next, we detected the protrusion onset and aligned the time-series protrusion velocities as a temporal fiduciary (**Fig. 1b.3-4**)^5^. Following the same procedure in our previous study^6^, we denoised the time series velocity profile using Empirical Mode Decomposition (EMD)^41^ to reduce noise.

To develop an unsupervised teach model (**Fig. 1c**), we employed auto-correlation functions (ACFs) as time series features (ACF-based clustering, **Fig. 1c. 2**). In our earlier study, ACFs produced highly interpretable protrusion phenotypes based on time series with uniform temporal lengths^6^. Nonetheless, subcellular protrusion occurs over various temporal durations, leading to heterogeneous time series lengths. Evaluating the similarity between samples of varying temporal lengths is inherently complex. Consequently, rather than computing the similarity distance across all samples, we specifically calculated distances among samples with comparable temporal lengths. To compute the ACF-based distance for time series with similar temporal lengths, we introduced random noises as padding to the latter part of the time series, thereby equalizing their temporal lengths (**Fig**. **1b****.5**). We followed our previous ML pipeline, which involved SAX (Symbolic Aggregate approXimation)^42^ (**Fig. 1c.1**) and Euclidean-based ACF distance. The distances, indicative of similarity, were aggregated to form a partial similarity distance matrix. Applying a community detection clustering algorithm^43^ to this matrix identified distinct clusters representing potential protrusion phenotypes (**Fig. 1c.3**). These clusters served as the basis for generating pseudo-labels crucial for training a student DNN. Additionally, we quantified the effects of molecular perturbations within each cluster. This quantification identified the clusters whose sensitivity to molecular perturbations would be maximized during subsequent training of the student model.

We proceeded with feature learning through the training of a student DNN model (**Fig. 1d**) utilizing the pseudo-labels from the teacher model. While standard classification tasks enable DNNs to learn crucial features discriminating existing labels, they may not adequately preserve the heterogeneity vital to deep phenotyping. Although autoencoder^20^ and its variants^44–47^ have been widely used for feature learning by minimizing the reconstruction loss, their features are not guaranteed to be discriminative to molecular perturbations of interest, and may contain irrelevant heterogeneity. To address these challenges, we introduced a novel regularizer, named SENSER (SENSitivity-enhancing autoEncoding Regularizer). Importantly, regularization plays pivotal roles in self-training^48,49^. By injecting significant heterogeneity during this feature learning, SENSER allows our student model to explore the feature space where specific molecular perturbations can be discriminated. To accomplish this, the student model training is regularized to maximize the effect size of specific molecular perturbations within the clusters identified by the teacher model. This novel regularization strategy is designed to capture features that preserve the essential heterogeneity in the context of molecular perturbations. Therefore, this approach markedly differs from supervised autoencoder^47^, where the primary focus is maximizing the accuracy of a given classification task. Specifically, we produce distinct clustering outcomes by adjusting the regularization parameter assigned to the losses of the autoencoder and the classifier. Subsequently, we compute the effect sizes of the clusters identified by the teacher model as sensitive to molecular perturbation, determining the optimal regularization parameter by maximizing these effect sizes. Through this process, our student model guarantees that the learned features encapsulate essential heterogeneity and exhibit discriminative power concerning specific molecular perturbation. Crucially, our method can also be viewed as a multi-task learning (MTL) approach, anticipated to yield more robust and generalizable features compared to single-task learning^39,47,50^.

Our student model was based on bidirectional Long-Short Term Memory (Bi-LSTM) structure^51,52^ (**Fig. 1d**). In particular, LSTM specializing in time series data can handle variable-length time series since it does not require a fixed length of time series. Having initially preprocessed the time series by rescaling them to the range [-1, 1] to mitigate the impact of large velocity magnitude (**Fig. 1b.6**), we trained our student model using our dataset, including more than 30,000 time-series samples. By maximizing the total loss comprising the loss functions of the autoencoder and the classifier, we extracted ‘deep features’ (features learned from deep learning). Subsequently, we applied Principal Component Analysis (PCA) to reduce the feature dimension and selected the first 15 principal components, explaining more than 95% of the total variance. Following this, we conducted clustering analysis to refine the preliminary clustering results obtained from the teacher model (**Fig. 1e**).

The deep features derived from our student model are anticipated to maintain the heterogeneity of molecular perturbation effects owing to the SENSER. Leveraging these deep features, we conduct sub-clustering analyses to further partition the identified clusters (parent phenotypes). We analyze the sub-clusters under control and perturbed conditions to identify those with heightened responsiveness to molecular perturbations. These specific sub-clusters are designated as ‘deep phenotypes’ (**Fig. 1e**). Subsequently, we utilize the proportions of subcellular protrusion phenotypes in individual cells as single-cell features for clustering analysis, facilitating the identification of single-cell protrusion phenotypes.

The interpretation of the identified deep phenotypes involves a two-step approach (**Fig. 1f**). First, statistical analyses are applied to ascertain the significance of deep features associated with identified deep phenotypes. Following this, SAGE (Shapley Additive Global ImportancE) analysis^53^ is employed to identify the temporal intervals that significantly contribute to the expression of these deep features. SAGE assigns values to each time point, elucidating their importance in predicting target feature values. This combined approach not only validates the relevance of identified features but also provides a global understanding of the temporal dynamics, pinpointing specific time points or durations crucial for shaping the deep phenotypes.

### Unsupervised teacher model with interpretable features for cell protrusions

Our unsupervised teacher model builds on our previous study^6^, where we utilized ACF features of the time series data and the density-peak clustering algorithm to cluster protrusion velocity time-series^54^. The previous analysis resulted in a poorly characterized heterogeneous cluster of “fluctuating” protrusion and was limited by the constraint of utilizing time-series of uniform temporal lengths for clustering. To address these limitations, we employed the community detection method^55^ to uncover more refined protrusion phenotypes. Additionally, this approach enables the utilization of all available data, irrespective of the lengths of the time-series, during the training of the student model, thereby aiding in the prevention of overfitting.

First, to know the role of the community detection method in our analysis, we focused on the equal length time series (56 time frames) of the protrusion velocity. By quantifying Silhouette values^56^ with varying the number of clusters (**Supplementary** Fig. 1a), we chose the optimal number of clusters as six. **Fig. 2a** shows the average velocity profiles in each cluster. The raw velocity map in each cluster (**Fig.2b, Supplementary** Fig. 1d), the t-SNE, and the ordered distance map of clustering results (**Supplementary** Fig. 1b**-c**) visually confirmed the distinctiveness of the clusters. The significant difference from our previous results was that we were able to split the previous “fluctuation” cluster into the clusters of “steady” (Cluster I) and “bursting” (Cluster II) protrusions (**Fig. 2a-b**). Because Cluster II exhibited rapid changes of edge velocity within 100 seconds, we named it “bursting protrusion”. Since Cluster III-V exhibited periodic edge velocity, we named them “periodic protrusion” and Cluster VI named “accelerating protrusion” as we did in our previous work^6^.

**Figure 2.**
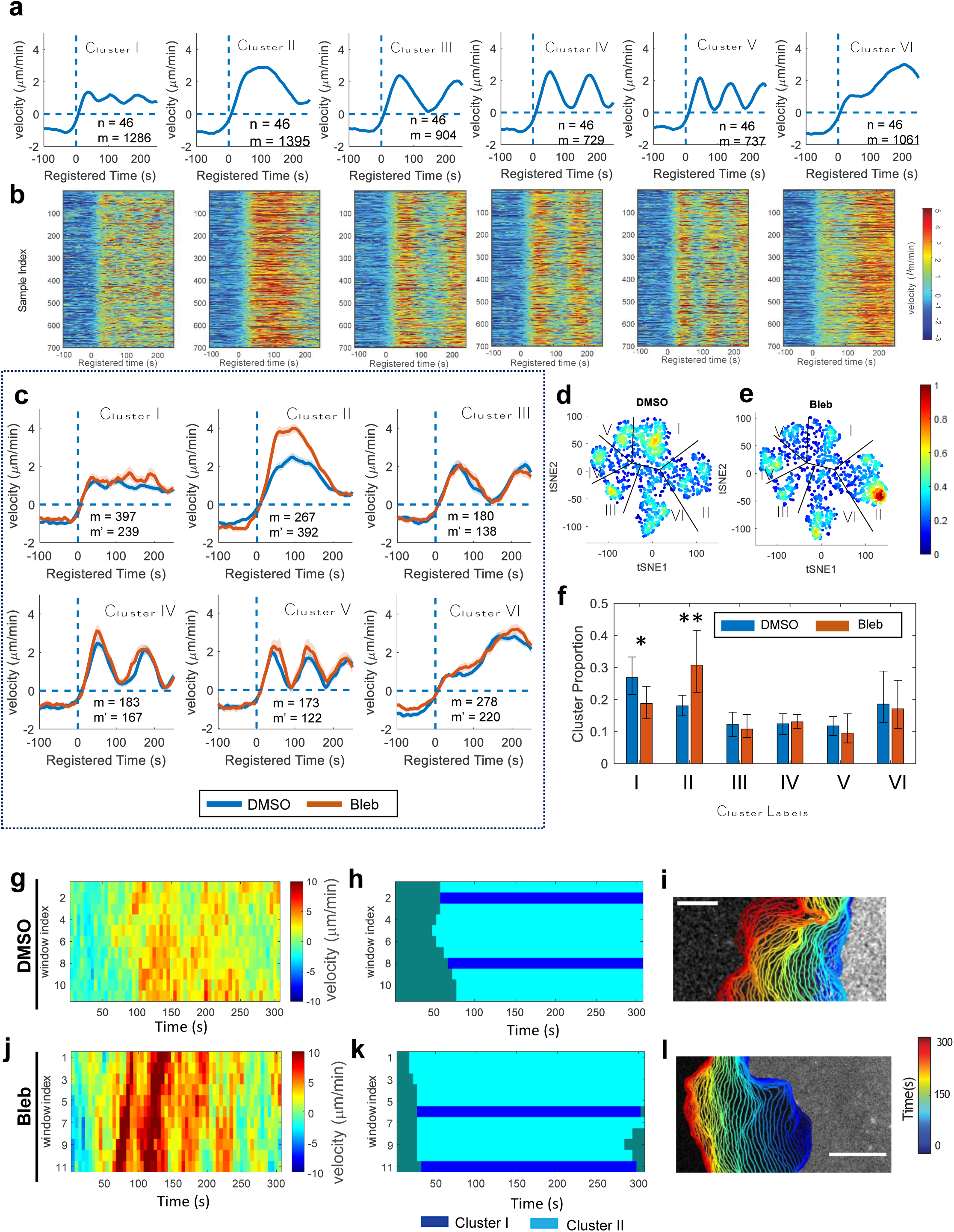
Pre-identification of subcellular protrusion phenotypes. (**a**) Average temporal patterns of protrusion time series in the clusters whose temporal length is larger than 250s with a 95% confidence interval registered at protrusion onset (*t* = 0). (m: the number of probing windows, the number of cells: 46). (**b**) Raw velocity maps for six identified phenotypes. All time series are registered with respect to protrusion onset (t = 0) (**c-f**) Effects of myosin-II inhibitor, blebbistatin on the subcellular protrusion phenotypes. Averaged protrusion velocity time series in each cluster registered at protrusion onset in DMSO or blebbistatin-treated cells (m: the number of probing windows in DMSO; m’: the number of probing windows in blebbistatin-treated cells) (c). t-SNE plots of autocorrelation functions of protrusion velocity time series overlaid with the density of data in the cells treated with DMSO (the number of cells: 14) (d) and blebbistatin (the number of cells: 13) (e). Comparison of the proportion for each cluster per cell between DMSO and blebbistatin (20 μM) (f). Solid lines indicate population averages, and shaded error bands indicate 95% confidence intervals of the mean computed by bootstrap sampling. The error bars indicate 95% confidence interval of the mean of the cluster proportions. * p<0.05, and ** p < 0.01 indicate the statistical significance by bootstrap sampling. (**g-l**) Representative bursting protrusion (Cluster II) in DMSO (g-i) and blebbistatin (j-l) treated cells. Raw velocity maps (g, j). Cluster labels (h, k). Edge time-evolution (I, l). Bars: 5 μm.

The well-established mechanism of cell protrusion at the leading edge is primarily driven by rapid actin polymerization, a process tightly regulated by various actin regulators, including Arp2/3 and VASP^57–60^. The specific temporal coordination between Arp2/3 and VASP has been characterized, particularly in the context of accelerating protrusion^6^. Additionally, myosin II plays a multifaceted role in regulating actin dynamics^61^, contractility^62–64^, and adhesion dynamics^65,66^, all of which contribute to the modulation of cell protrusion. To discern the specific effects of these regulators in each identified phenotype, we conducted an analysis of live cell movies from drug-treated experiments. Initially, we performed the analysis of the CK666^67^ (Arp2/3 inhibitor) perturbation to these protrusion phenotypes. Consistently with the previous results, CK666 significantly reduced the proportion of the accelerating protrusion phenotype (Cluster VI) only in comparison to the inactive control (CK689) (p-value=0.013 by bootstrap resampling) (**Supplementary** Fig. 1e-h).

Next, we characterized the effects of blebbistatin on each phenotype. First, the velocity profile in Cluster II (bursting protrusion) was substantially elevated by blebbistatin treatment (**Fig. 2c**) and **Fig. 2g-l** demonstrate an example of bursting protrusion of a blebbistatin-treated PtK1 cell in comparison to that of the control case. Moreover, the t-SNE visualization revealed that more bursting protrusions (Cluster II) were produced by blebbistatin treatment (**Fig. 2d-e**). Consistently, the quantification of the proportion of the clusters showed that bursting protrusion was significantly increased by the blebbistatin treatment (p-value=0.004 by bootstrap resampling) (**Fig. 2f**), while CK666 did not show significant effects (**Supplementary** Fig. 1h). This result aligns with a previous study suggesting that downregulation of myosin II with blebbistatin promotes lamellipodial protrusion^65^. Nevertheless, our analysis implies that this form of protrusion can coexist with a range of diverse activities within the spectrum of protrusion behaviors.

As distinct subcellular protrusion phenotypes may arise due to diverse spatiotemporal regulation of actin regulators, we investigated the relationship between the velocity profiles of each protrusion phenotype and the dynamics of fluorescence intensities in actin and several actin regulators associated with each protrusion phenotype. This analysis was conducted using time-lapse movies of PtK1 cells expressing fluorescently tagged actin (SNAP-tag-actin), Arp3 (HaloTag-Arp3), and VASP (HaloTag-VASP), with a cytoplasmic marker (HaloTag) serving as a control signal. The time-series fluorescence intensities of each molecule were averaged for each protrusion phenotype (**Supplementary** Fig. 2). While accelerating protrusion (Cluster VI) exhibited the same dynamics of actin and actin nucleators as our previous study^6^, the fluorescence intensities of actin and VASP in Cluster IV-V exhibited much clearer periodicity (**Supplementary** Fig. 2a and c) than the previous study where their intensity periodicity was lost as the frequency of the oscillation increased (Fig. 3 in the previous study^6^). Intriguingly, the intensities of Arp3 still did not show periodicity in Cluster IV-V (**Supplementary** Fig. 2b), suggesting that Arp2/3 and VASP may have differential roles in regulating Periodic Protrusions.

**Figure 3.**
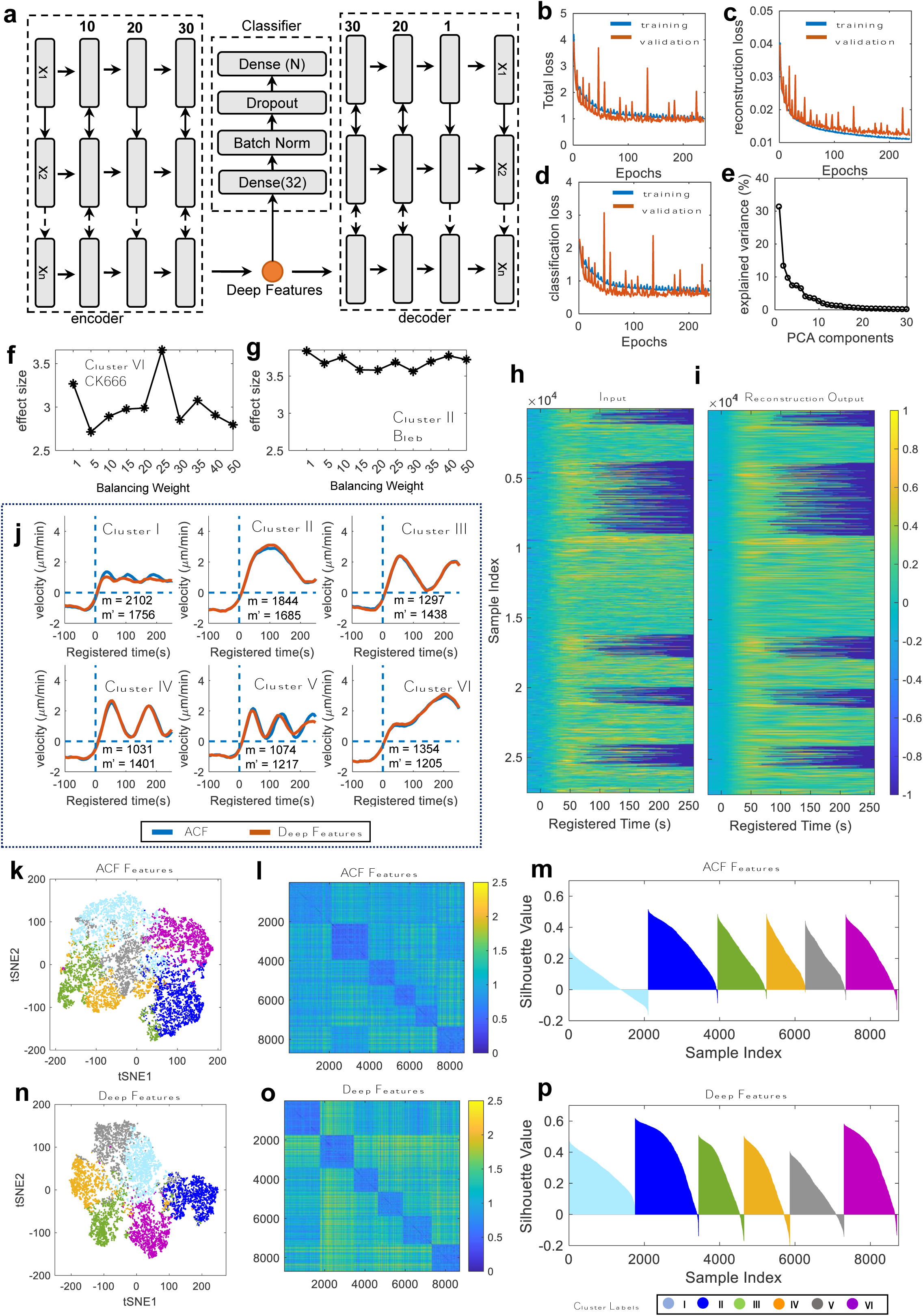
Deep feature learning from protrusion velocity time-series. (**a**) Deep feature learning of supervised bi-LSTM autoencoder. (**b-d**) The training performance of feature learning. Total loss (b), autoencoder loss (c) and classification loss (d). (**e**) Principal component analysis on the deep features. (**f**) Effect sizes of the difference between CK689 and CK666 in Cluster VI with varying weight parameters. (**g**) Effect sizes of the difference between DMSO and blebbistatin in Cluster II with varying weight parameters. (**h-i**) Visual comparison between the input of scaled velocities (h) and reconstructed output from autoencoder (i). (**j-p**) Comparison between ACF and DF-based clustering. Average protrusion velocity time series registered at protrusion onset (t=0) (j). Solid lines indicate population averages, and shaded error bands indicate 95% confidence intervals of the mean computed by bootstrap sampling. (the number of cells: 81; m: the number of probing windows in ACF-based clustering; m’: the number of probing windows in Deep Feature-based clustering). t-SNE plot of ACF (k) and DF (n). Ordered distance map from ACF-based clustering (i) and DF-based clustering (o). Silhouette plots from ACF-based clustering (m) and DF-based clustering (p).

After confirming the beneficial effects of the community detection algorithm in our teacher model, we added the time series of variable temporal lengths to the dataset and performed the same clustering analysis using the partial similarity matrix of ACF distances (**Supplementary** Fig. 3a-b). The clustering results contained the same types of the clusters (Cluster I∼VI) as the equal length time series. We also had additional clusters whose temporal lengths are shorter than 250s (Cluster VII∼X). Cluster VII and VIII are short (< 200s) lived transient protrusions and Cluster IX and X are slightly longer (>200s) protrusions. In summary, our findings suggest that using the community detection algorithm in this study helped to generate more accurate labels for time-series data of varying lengths, which will improve the training of our student model.

### Deep feature learning for subcellular protrusion phenotyping

We used pseudo-labels created by the teacher model for time-series data of varying temporal lengths to train our student DNN (**Fig. 3a**). As training progressed, we observed a decrease in the reconstruction, classification, and total losses (**Fig. 3b-d**). The visual comparison between the input and the output confirmed that the training was effective (**Fig. 3h-i**). After reducing the dimensionality of the biLSTM encoder features (Deep Features, DF) by PCA (**Fig. 3e**), we applied the community detection algorithm to identify clusters. To determine the regularization parameter of SENSER that balances the losses of the autoencoder and the classifier, we evaluated the clustering results based on the effect sizes of accelerating protrusion (Cluster VI) between CK689 and CK666 (**Fig. 3f**), and bursting protrusion (Cluster II) between DMSO and blebbistatin (**Fig. 3g**). We found that the effect size of accelerating protrusion was highest when the regularization parameter was set to 25 (**Fig. 3f**), while there was no clear pattern for the effect size variations of bursting protrusion (**Fig. 3g**). This means that finding the optimal regularization parameter is critical for the analysis of accelerating protrusion, while the wide range of the regularization parameters can be used for the analysis of bursting protrusion. Hence, we determined the optimal regularization parameter for the student model as 25.

We observed that the average patterns of protrusion velocities resulting from clustering based on both ACF and DF, with equal temporal lengths, exhibit similarities (**Fig. 3j**) (**Supplementary** Fig. 3c-d illustrates the results for variable temporal lengths). However, the t-SNE feature visualization highlighted notable distinctions between the two methods, revealing clearer cluster boundaries when employing DF (**Fig. 3k** and **n**). The utilization of DF for clustering enhanced the clustering outcomes, providing a distinct order-distance map and yielding high silhouette values (**Fig. 3l**, **m**, **o**, and **p**). This enhancement, facilitating a more robust and detailed analysis of each protrusion type, can be attributed to our self-training strategy.

To evaluate the model’s generalization to a different cell line, we applied the student model trained on PtK1 cells to analyze protrusion velocity time series of MCF10A, human mammary epithelial cells. Clustering results using these DFs demonstrated that MCF10A cells exhibit bursting and accelerating protrusion phenotypes, in addition to steady and periodic phenotypes (**Supplementary** Fig. 4). This suggests that our findings extend to other types of epithelial cell lines.

### Deep features enhance the sensitivity of statistical analysis

In our CK666 perturbation analysis, where we optimized the deep feature learning, we found similar clustering patterns using the DFs (**Fig. 4a**) as those obtained using the ACF (**Supplementary** Fig. 1e). We observed that the significant drug effects of CK666 on Clusters I and VI (**Fig. 4b-c**) are consistent with the results from ACF-based teacher model (**Supplementary** Fig. 1f-h). While both ACF-based and DF-based clustering showed significant results (ACF-based clustering, p-value=0.045 for Cluster I and 0.013 for Cluster VI; DF-based clustering, p-value = 0.016 for Cluster I and 0.0069 for Cluster VI by bootstrap resampling), we found that DF-based clustering showed smaller p-values, demonstrating that the DF can provide more sensitive and robust statistical analyses.

**Figure 4.**
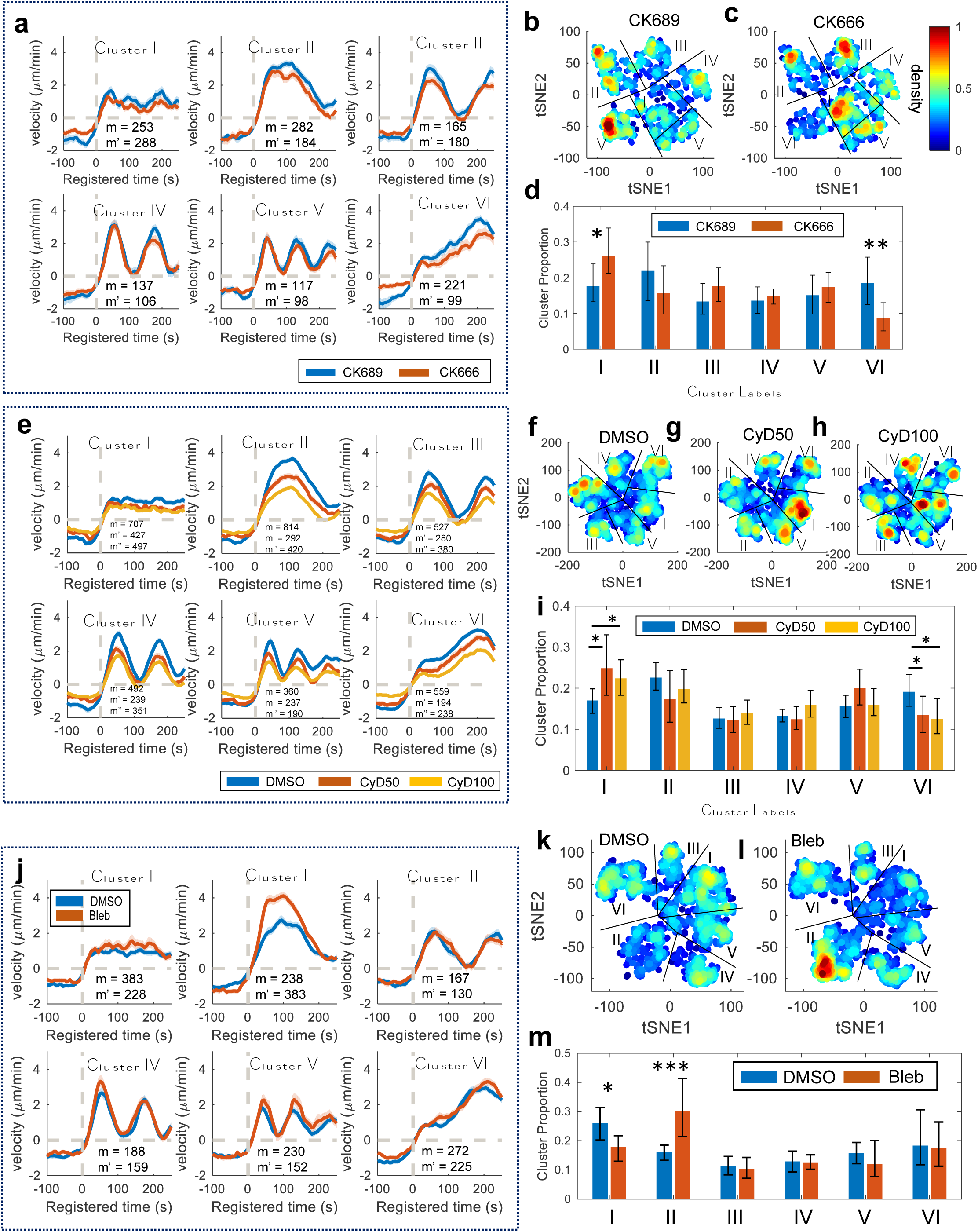
Effects of CK666 and Cytochalasin D on subcellular protrusion phenotypes. (**a, e**) Average velocity time series registered at protrusion onset (t=0) in each cluster of the cells treated with CK689 or CK666 (m: the number of probing windows in CK689(control)-treated cells; m’: the number of probing windows in CK666-treated cells) (a) and DMSO or Cytochalasin D (CyD50: 50 nM, CyD100: 100nM) (m: the number of probing windows in DMSO-treated cells; m’: the number of probing windows in CyD50-treated cells; m’’: the number of probing windows in CyD100-treated cells) (e). Solid lines indicate population averages, and shaded error bands indicate 95% confidence intervals of the mean computed by bootstrap sampling. (**b-c, f-h**) t-SNE plot overlaid with the density of the deep features of the cells treated with CK689 or CK666 (b-c) and DMSO or Cytochalasin D (f-h). (**d, i**) Comparison of the proportion for each cluster per cell between CK689 and CK666 (d), and DMSO and Cytochalasin D (i). (**j**) Average velocity time series registered at protrusion onset (t=0) in DF-based clusters of the cells treated with DMSO or blebbistatin (m: the number of probing windows in DMSO; m’: the number of probing windows in blebbistatin-treated cells). (**k-l**) t-SNE plot overlaid with the density of the deep features of the cells treated with DMSO (k) or blebbistatin (l). (**m**) Comparison of the proportion for each cluster per cell between DMSO and blebbistatin. The error bars indicate a 95% confidence interval of the mean of the cluster proportions. *p<0.05, **p<0.01 indicate the statistical significance by bootstrap sampling. The numbers of cells: 10 for CK689, 10 for CK666 (a-d) and 22 for DMSO, 16 for CyD50, 20 for CyD100 (e-i).

To confirm the validity of our results, we used the same DF-based clustering on two additional drugs: Cytochalasin D (CyD) at concentrations of 50-100nM (**Fig. 4e-i**) and blebbistatin (Bleb) (**Fig. 4j-m**). Previous research has shown that CyD affects accelerating protrusion by inhibiting VASP-dependent actin elongation^6^. We found that CyD significantly decreased the proportion of Cluster VI (CyD at 50nM: p-value = 0.027; CyD at 100nM: p-value = 0.013 by bootstrap resampling) (**Fig. 4i**), and significantly increased the proportion of Cluster I (CyD at 50nM: p-value = 0.021; CyD100 at 100nM: p-value = 0.023 by bootstrap resampling), consistent with the previous finding using CK666. Additionally, blebbistatin significantly elevated the proportion of Cluster II (p-value = 0.0004; by bootstrap resampling), surpassing the p-value obtained using the teacher model with ACF-based clustering (p-value=0.004, **Fig. 4j, m**). In contrast, Cluster II was not significantly affected by CK666 (**Fig. 4d**, p-value = 0.4577; bootstrap sampling) or CyD (**Fig. 4i**, CyD50: p-value = 0.0707, CyD100: p-value = 0.1455; bootstrap sampling). The t-SNE plot of the DF distribution displays that the local region of Cluster II was highly elevated by the blebbistatin treatment (**Fig. 4k,i**).

### Deep phenotypes of accelerating and bursting protrusions

With the validation of the enhanced statistical analysis provided by DeepHACX’s deep features for parent phenotypes, our focus shifts to the exploration of hitherto undiscovered deep phenotypes—subtypes within the broader parent categories—and their associated drug sensitivities. Our investigation commences with a detailed sub-clustering analysis, concentrating initially on samples within Cluster VI, characterized as accelerating protrusions. The aim is to pinpoint fine-grained clusters within this category that exhibit heightened sensitivity to drug perturbations. The t-SNE analysis of DFs suggested the presence of sub-clusters that may be more sensitive to CK666 (**Fig. 5a**) and Cytochalasin D (**Fig. 5b**). The Silhouette values indicated that the optimal number of sub-clusters of accelerating protrusion was three (**Supplementary** Fig. 5a-b). The sub-clusters identified in Cluster VI displayed subtle but distinct behaviors (**Fig. 5c, e**, and **6c**, **Supplementary** Fig. 5c). We identified one weak (Cluster VI-1) and two strong accelerating clusters (Cluster VI-2 and 3). Cluster VI-2 exhibited a brief pause in acceleration before 100s, while Cluster VI-3 had constant acceleration.

**Figure 5.**
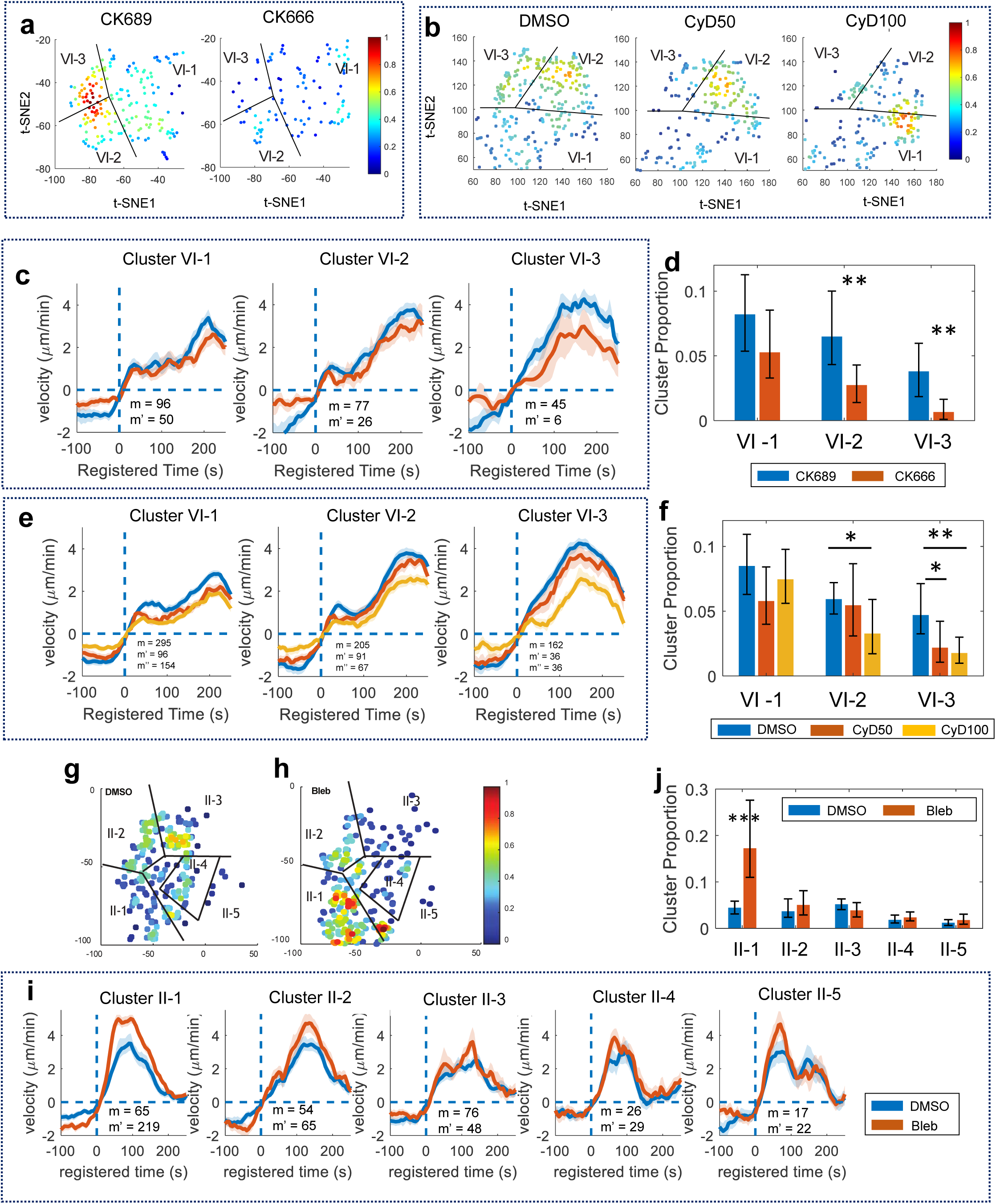
Deep phenotypes in accelerating protrusion and bursting protrusion. (**a-b**) t-SNE plots overlaid with the density of the deep features from accelerating protrusion (Cluster VI) for CK689/CK666-treated (a) or DMSO/Cytochalasin D treated cells (b). (**c-d**) Average protrusion velocity time series (m: the number of probing windows) (c) registered at protrusion onset (t=0) and comparison of the proportion (d) for three deep phenotypes of accelerating protrusion from the cells treated with CK689 or CK666 (m: the number of probing windows in CK689-treated cells; m’: the number of probing windows in CK666-treated cells). (**e-f**) Average velocity time series registered at protrusion onset (t=0) (e) and comparison of the proportion (f) of the deep phenotypes of accelerating protrusion from the cells treated with DMSO/Cytochalasin D (m: the number of probing windows in DMSO-treated cells; m’: the number of probing windows in CyD50-treated cells; m’’: the number of probing windows in CyD100-treated cells). (**g-h**): t-SNE plots overlaid with the density of the deep features from the Bursting protrusion (Cluster II) for DMSO (g) or blebbistatin treated cells (h). (**i-j**): Average velocity time series registered at protrusion onset (t=0) (i) and comparison of the proportion (j) of the deep phenotypes of Bursting protrusion from the cells treated with blebbistatin. (m: the number of probing windows in DMSO-treated cells; m’: the number of probing windows in blebbistatin - treated cells). Solid lines indicate population averages, and shaded error bands indicate 95% confidence intervals of the mean estimated by bootstrap sampling. The error bars indicate a 95% confidence interval of the mean of the cluster proportions. *p < 0.05, **p<0.01 indicate the statistical significance by bootstrap sampling. The numbers of cells: (c-d) CK689: 10, CK666: 10; (e-f) DMSO: 22, CyD50: 16, CyD100: 20; (i-j) DMSO: 14, blebbistatin: 13.

Then, we found that Cluster VI-1 was not significantly affected by CK666 (**Fig. 5c-d**; p-value: 0.0694, bootstrap resampling) and low doses of CyD (**Fig. 5e-f**; CyD at 50nM: p-value = 0.0505; CyD at 100nM: p-value = 0.2543, bootstrap resampling). However, the proportions of Cluster VI-2 and 3 were significantly decreased by CK666 (**Fig. 5d**; p-value = 0.0049 for VI-2, 0.0012 for VI-3, bootstrap resampling) and CyD (**Fig. 5f**; CyD at 50nM: p-value = 0.3698 for VI-2, 0.0193 for VI-3; CyD at 100nM: p-value = 0.0165 for VI-2, 0.0011 for VI-3, bootstrap resampling). Furthermore, the effects of CK666 and CyD on Cluster VI-3 were stronger than Cluster VI-2. The treatment with blebbistatin had no significant effect on any deep phenotypes of accelerating protrusion (**Supplementary** Fig. 5d-e). We also identified similar deep accelerating phenotypes from MCF10A cells by using the DFs from the student model trained with PtK1 cells (**Supplementary** Fig. 5f-g). In particular, Cluster VI-3 from MCF10A exhibited a more robust acceleration profile than the other subclusters, sharing the same characteristics as PtK1 cells.

We conducted a parallel sub-clustering analysis on Cluster II (bursting protrusions) to further identify deep phenotypes sensitive to blebbistatin. Sub-clustering utilizing DFs identified five clusters based on the silhouette values (**Supplementary** Fig. 6a-b). These deep phenotypes (Cluster II-1∼5) identified in Cluster II were shown displayed subtle and visually distinct behaviors as shown in **Fig. 5i** and **6f** and **Supplementary** Fig. 6c. We found that Cluster II-1 was significantly increased by blebbistatin (p-value < 0.0001, bootstrap sampling) while the other Clusters (II-2∼5) were not significantly affected (**Fig. 5g-j**). This refined deep phenotyping pinpointed Cluster II-1 as the sole phenotype susceptible to blebbistatin treatment. These findings underscore the utility of our deep phenotyping approach in isolating specific phenotypes, offering precise quantification of drug actions.

**Figure 6.**
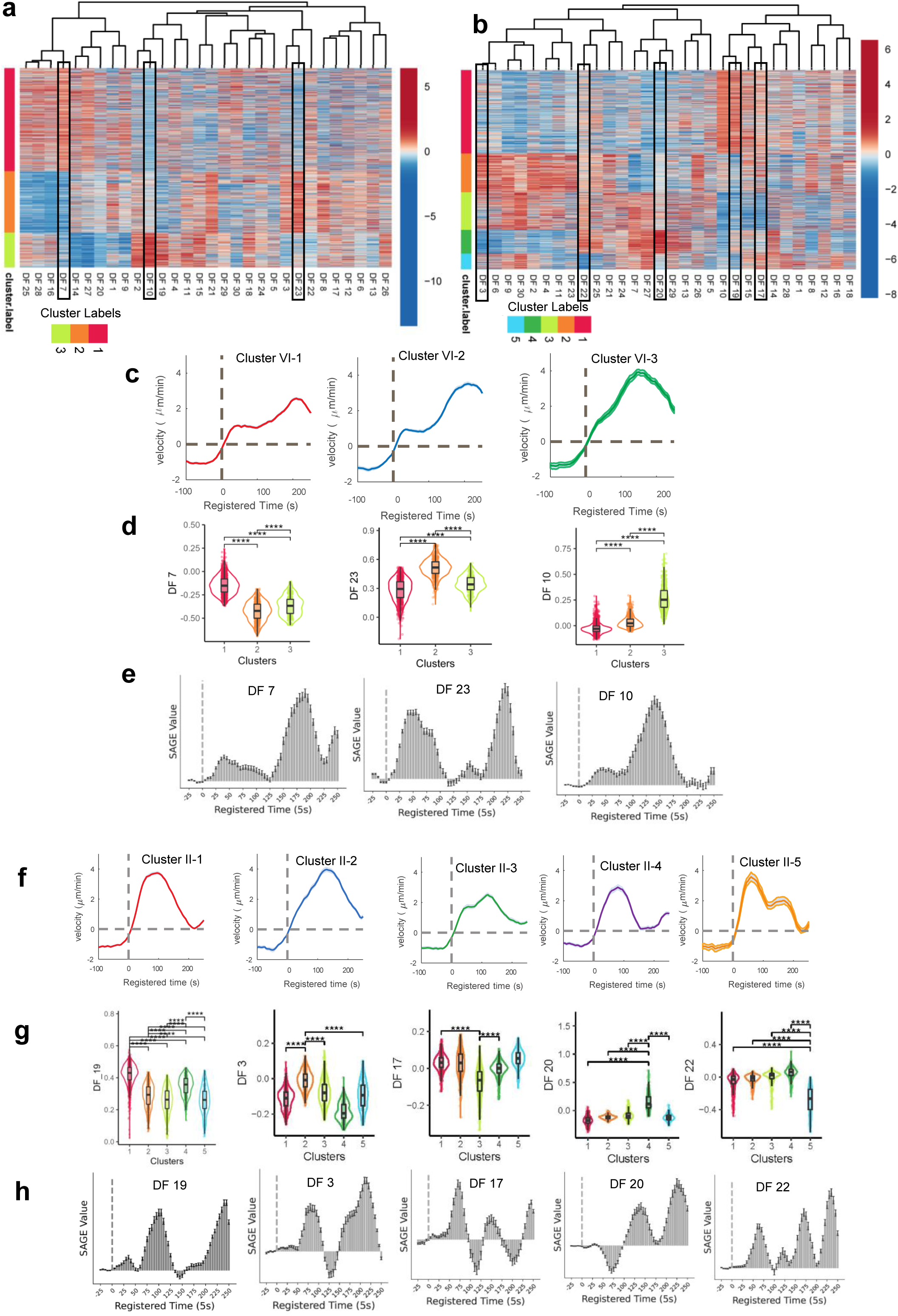
Interpretation of deep features associated with deep phenotypes. (**a-b**) Visualization of deep features in the phenotypes of Accelerating (a) and bursting protrusion (b). The rectangular boxes indicate the deep features uniquely linked to deep phenotypes. (**c**, **f**) Average velocity time series of deep phenotypes of Accelerating (c) and Bursting (f) Protrusion registered at protrusion onset (t=0) (m: the number of probing windows). (**d**, **g**) Differential expression analysis of deep features with Accelerating (d) and Bursting (g) Protrusion. **** indicates statistical significance with p<0.0001. The p values were estimated using the Wilcoxon rank sum test implemented in *presto* R package. (**e**, **h**) SAGE analysis of each deep feature in Accelerating (e) and Bursting (h) Protrusion.

In contrast, CK666 treatment did not affect any deep phenotypes of Cluster II significantly (**Supplementary** Fig. 6d-e and **i,k**). CyD treatment decreased the proportion of Cluster II-2 and 5 significantly (p-value < 0.0001 for II-2 and CyD100, p-value = 0.0059 for II-5 and CyD50) and increased the proportion of Cluster II-3 and 4 significantly (p-value = 0.004 for Cluster II-3 and CyD100, p-value = 0.0149 for Cluster II-4 and CyD100) (**Supplementary** Fig. 6f-h and **j,l**). Due to the opposite effects of CyD on the deep phenotypes of busting protrusion, the original bursting phenotype (Cluster II) was not shown to be significantly affected by CyD (**Fig. 4i**). We also identified bursting deep phenotypes from MCF10A cells by using the DFs from the student model trained with PtK1 cells (**Supplementary** Fig. 6m-n). Clusters II-1∼3 in MCF10A cells displayed a velocity profile remarkably similar to that of PtK1 cells, whereas Clusters II-4∼5 exhibited distinctive differences from PtK1 cells. Collectively, our deep phenotyping approach facilitated the identification of nuanced temporal patterns within our protrusion velocity time series dataset, intricately associated with specific drug perturbations.

### Interpretation of deep phenotypes

We employed a comprehensive interpretation approach to elucidate the distinct roles played by deep features in various deep phenotypes identified by DeepHACX. Visualizations of the DF values (without PCA), organized for acceleration protrusion (**Fig. 6a**) and bursting protrusion (**Fig. 6b**) phenotypes, highlighted their individual contributions to each deep phenotype. In the case of accelerating protrusion, the elevated values of DF7, DF23, and DF10 were significantly associated with Clusters VI-1, 2, and 3, respectively (**Fig. 6d**, Wilcoxon rank sum testing). For bursting protrusion, the heightened values of DF19, DF3, and DF20 were distinctly linked with Clusters II-1, 2, and 4, respectively, while the diminished values of DF17 and DF22 were associated with Clusters II-3 and 5, respectively (**Fig. 6g**, Wilcoxon rank sum testing). The presence of DFs exclusively correlated with each deep phenotype underscores the remarkable efficiency of DeepHACX’s features. With a minimal number of DFs, DeepHACX can comprehensively elucidate the heterogeneity characterizing both accelerating and bursting protrusion phenotypes.

The identification of a concise set of DFs, each exclusively associated with a specific deep phenotype, forms the basis for the subsequent SAGE analysis^53^. This interpretability analysis unveils the individual time points’ contribution to predicting distinct deep phenotypes. As each deep phenotype aligns exclusively with a particular DF, capturing essential heterogeneity, discerning each time point’s contribution to a specific DF enhances the interpretability of the associated deep phenotype. Thus, the SAGE values for each DF provide insights into crucial time intervals for the manifestation of each deep phenotype. The deep phenotypes of accelerating protrusion displayed highly specific temporal contributions (**Fig. 6c** and **e**). The importance of protrusion events in Cluster VI-3 is emphasized by the SAGE values associated with DF10, particularly highlighting events occurring between 100 and 180s. Conversely, in Cluster VI-2, the crucial time points are those before 100s and after 180s, as indicated by the SAGE values linked to DF23. Similarly, for Cluster VI-1, DF7’s SAGE values highlight the significance of protrusion events occurring after 150s.

In addition, the SAGE analysis for the deep phenotypes of bursting protrusion revealed precise temporal contributions (**Fig. 6f** and **h**). Despite the similar dynamic profiles observed in Clusters II-1, 2, and 4, the SAGE profiles exhibited remarkable specificity for each cluster. In Cluster II-1 (DF19), pivotal protrusion events were identified around 100s when the protrusion velocity peaked and after 200s when protrusion ceased. Conversely, in Cluster II-2 (DF3), crucial protrusion events occurred around 80s, during a phase of increasing protrusion velocity. Notably, time points around 125s, when protrusion velocity reached its maximum, did not significantly contribute to Cluster II-2. For Cluster II-4 (DF20), the significant temporal interval was after 125s, representing the late phase of protrusion. This underscores the distinct and specific occurrence of significant events in these deep phenotypes of bursting protrusion at precise time intervals. Notably, in Clusters II-3 (DF17) and 5 (DF22), as suggested by their velocity profiles (**Fig. 6f**), multiple time intervals contribute to their phenotypes.

Our interpretation approach unraveled the remarkable properties of DeepHACX’s features, showcasing their distinctive efficiency in characterizing the heterogeneity within accelerating and bursting protrusion phenotypes. Through visualizations and statistical analyses of DFs, we identified a minimal set of features exclusively correlated with each deep phenotype. This precision laid the foundation for subsequent SAGE analysis, offering insights into the temporal intervals crucial for the manifestation of each fine-grained phenotype. The interpretability of DeepHACX’s features stems from their ability to encapsulate essential heterogeneity, enabling the discernment of individual time points’ contributions to the prediction of specific deep phenotypes.

### Subcellular phenotypes as single-cell features

Our subcellular phenotyping approach yields valuable insights into the molecular mechanisms governing cellular protrusion at a subcellular level. Furthermore, by leveraging these subcellular phenotypes as features at the single-cell level, we can unravel a deeper understanding of individual cell behaviors through a bottom-up analytical approach. The inherently interpretable nature of our subcellular protrusion phenotypes enhances the comprehensibility of single-cell phenotypes and their intricate relationship with underlying molecular mechanisms. To delineate single-cell phenotypes related to cell protrusion, we quantified the proportions of subcellular protrusion phenotypes within individual cells, utilizing them as cellular features. Subsequently, we applied the manifold learning technique, UMAP (Uniform Manifold Approximation and Projection)^68^, to visualize and analyze these single-cell feature distributions. With the feature distribution of single-cell data in a lower-dimensional space, we conducted clustering analysis using a community detection method.

The silhouette plots with the varying number of clusters indicated that the optimal number of cell clusters is nine (**Supplementary** Fig. 7a). UMAP 2D visualization (**Fig. 7a**), the proportion plots of each cluster (**Fig. 7b**), the ordered-distance plot (**Fig. 7c**), and silhouette plot (**Supplementary** Fig. 7b) demonstrated that the identified cell clusters are highly distinct. Each cell cluster was characterized by the mean proportions of subcellular protrusion phenotypes (**Fig. 7d**). Particularly, Cell Cluster 7 (C7 in **Fig. 7d**) and Cell Cluster 9 (C9 in **Fig. 7d**) have high levels of Bursting and accelerating protrusion. Therefore, we call them Bursting Cells and Accelerating Cells, respectively. Cell Cluster 8 (C8 in **Fig. 7d**) or Cell Cluster 6 (C6 in **Fig. 7d**) have high or medium levels of both Bursting and accelerating protrusion. Therefore, we call them Strong or Mid-Bursting/Accelerating Cells. In **Supplementary Table 1**, we summarized the characteristics of these cell clusters and their phenotype names. **Fig. 7i-n** depict representative examples of Cell Cluster 7-9.

**Figure 7.**
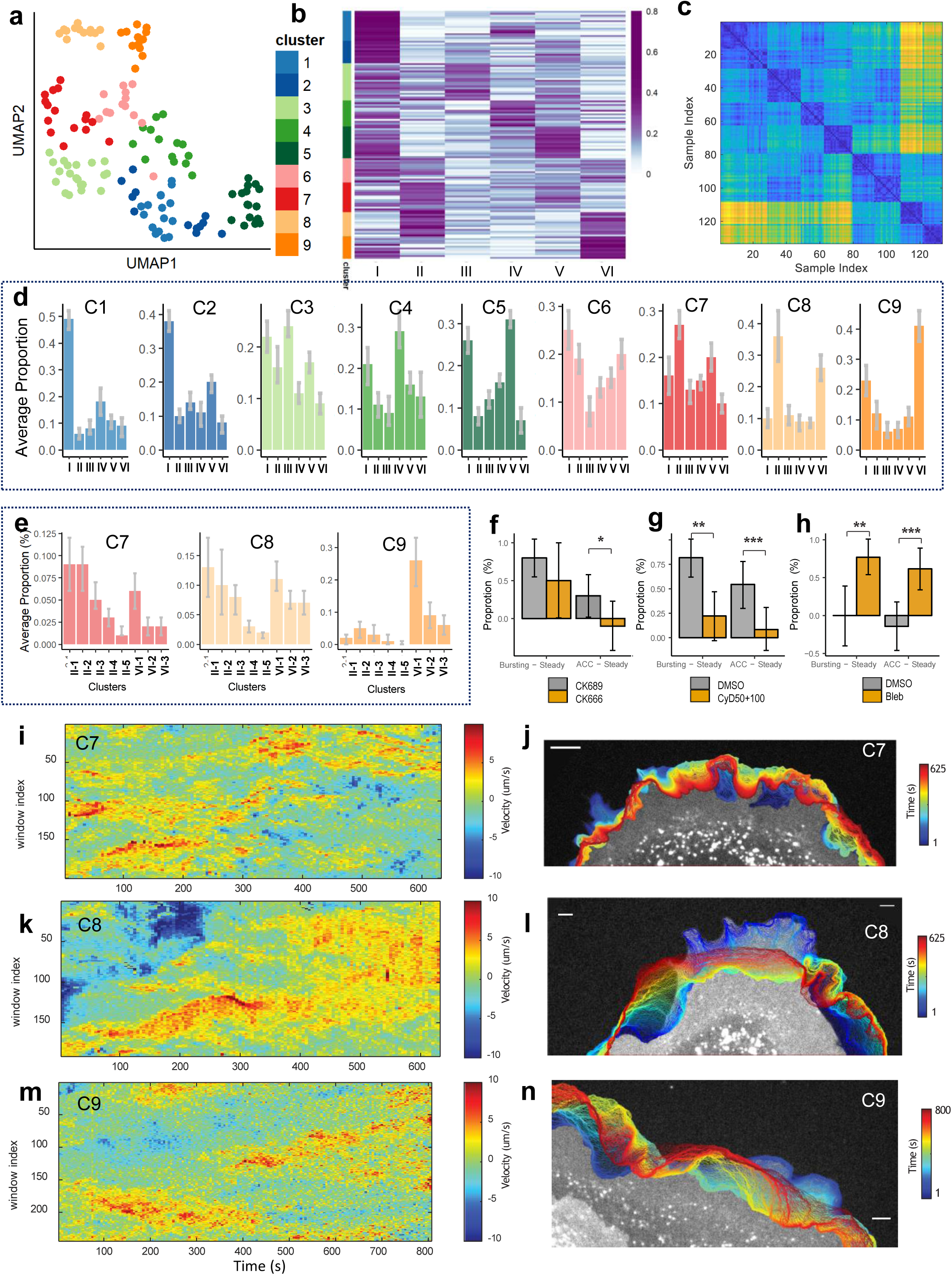
Single-cell phenotyping based on subcellular characterization. (**a-b**). UMAP (a) and heatmap (b) of clustered single-cell proportions of subcellular protrusion phenotypes (the number of cells: 133). (**c**) The ordered distance map of single-cell clustering. (**d-e**) Average subcellular protrusion proportions in each cell phenotype (d) and the deep phenotypes of bursting (Cluster II) and accelerating (Cluster VI) protrusions in Cell Phenotype 7∼9 (e). The 95% confidence intervals of the mean were estimated by bootstrap sampling. n: the number of cells in each cell cluster. (**f-h**) Comparison of the proportional differences between Bursting and Steady Cell Groups, and Accelerating and Steady Cell Groups in the conditions of CK689/CK666 (the number of cells in CK689-treated cells: 10, the number of cells in CK666-treated cells: 10) (f), DMSO/Cytochalasin D (the number of cells in DMSO cells: 22, the number of cells in Cytochalasin D-treated cells: 36) (g), DMSO/blebbistatin (the number of cells in DMSO cells: 14, the number of cells in blebbistatin-treated cells: 13) (h). *p<0.05, ***p<0.001 indicate the statistical significance by bootstrap sampling. (**i-n**) Representative examples of the space-time maps of instantaneous edge velocity of the cell edge of Cell Phenotype 7 (i), 8 (k), and 9 (m). Representative examples of edge evolution on 5 s interval of Cell Phenotype 7 (l), 8 (l), and 9 (n).

We also quantified the proportions of the subcellular deep phenotypes, the blebbistatin-sensitive Cluster II-1 (Bursting-1) and CK666/CyD-sensitive Cluster VI-2/3 (Accelerating 2/3) in each cell cluster (**Fig. 7e, Supplementary** Fig. 7c). Bursting Cells (Cell Cluster 7) have high levels of Cluster II-1 and low levels of Cluster VI-2/3. Conversely, Accelerating Cells (Cell Cluster 9) have a low level of Cluster II-1 and high levels of Cluster VI-2/3. Intriguingly, Strong-Bursting/Accelerating Cells (Cell Cluster 8) have high levels of both Cluster II-1 and VI-2/3, while they have fewer Cluster VI-1 (CK666/CyD-insensitive) than Accelerating Cells. As summarized in **Supplementary Table 1**, as cells have more proportions of bursting or accelerating protrusion, they tend to have more corresponding drug-sensitive deep phenotypes. This suggests that the identified cell phenotypes have differential sensitivities to CK666, CyD, and blebbistatin. To confirm this, we first pooled the cell phenotypes as follows: Cell Cluster #3, 6, 7, and 8 into Bursting Cell Group; Cell Cluster #6, 8, and 9 into Accelerating Cell Group; Cell Cluster #1 and 2 into Steady Cell Group. Since Steady Protrusion was affected by the drugs oppositely to Bursting or accelerating protrusion in the previous analysis (**Fig. 4d, i,** and **m**), we quantified the proportional differences between the cells of Bursting and Steady Cell Groups or Accelerating and Steady Cell Groups. We found that CK666 significantly decreased the proportion of Accelerating Cell Group over Steady Cell Group while it did not significantly change that of Bursting Cell Group over Steady Cell Group (**Fig. 7f**). Intriguingly, CyD/blebbistatin significantly decreased/increased the proportions of both Bursting and Accelerating Cell Groups over Steady Cell Group (**Fig. 7g-h**) even though they did not affect Bursting/accelerating protrusion in the previous subcellular analysis where every cell phenotype was considered (**Fig. 4i and 6c**). This demonstrates that both subcellular and cellular phenotyping is necessary to fully understand the heterogeneity of cellular drug responses.

## Discussion

In this study, we developed DeepHACX, a deep learning-based unsupervised self-training framework tailored for the analysis of live cell movies. Our results demonstrate that DeepHACX excels at learning concise and interpretable features that effectively capture the intricate heterogeneity induced by molecular perturbations. This unique capability positions DeepHACX as a powerful tool for a comprehensive exploration of molecular mechanisms concealed within subcellular and cellular heterogeneity.

In DeepHACX, the process of unsupervised learning is involved with the transfer of knowledge from an interpretable, unsupervised teacher model to a student model, facilitating the extraction of features that are both robust and generalizable. The teacher-student framework implemented in DeepHACX acts as a guiding mechanism for the student model, allowing it to capture essential information from the interpretable features of the teacher model. This knowledge transfer process, coupled with the application of our specialized regularizer, SENSER, encourages the student model to maximize the heterogeneity associated with molecular perturbations. Moreover, the identification of a minimal set of features contributing to specific deep phenotypes, strongly supports the notion that these features achieved through DeepHACX are closely aligned with disentangled features in the context of deep learning^18^.

Feature disentanglement implies that each learned feature distinctly represents a particular aspect or factor within the data^18^. In this context, the deep features identified by DeepHACX are not only concise but also exhibit a specificity that aligns with the unique characteristics of the underlying cellular dynamics and molecular perturbation effects. The minimal set of the features extracted by DeepHACX are disentangled in a way that each feature corresponds to specific aspects or deep phenotypes of cell protrusion. The fact that these deep features are concise yet robust enough to capture the heterogeneity suggests a form of disentanglement where each feature isolates and represents a distinct and interpretable aspect of the complex and multifaceted cellular dynamics

The SAGE analysis conducted in our study provides a crucial layer of validation for the achieved feature disentanglement by DeepHACX. SAGE analysis delves into the temporal contributions of specific features to the overall prediction of deep phenotypes, essentially pinpointing which time points play pivotal roles in shaping each phenotype. The observation that DeepHACX’s minimal set of features exhibits highly specific temporal contributions to various deep phenotypes aligns with the essence of disentangled features. This specific temporal attribution implies that each feature distinctly contributes to the manifestation of different phenotypes, thereby reinforcing the notion that DeepHACX has effectively disentangled these features. The precision offered by SAGE analysis in identifying the temporal intervals associated with deep features further enhances the interpretability and specificity of these features, solidifying their disentangled nature and instilling an additional layer of confidence in the reliability and robustness of our findings. This intrinsic disentanglement facilitates a more interpretable and granular understanding of the intricate interplay between cellular behaviors and molecular perturbations, underscoring the efficacy of the DeepHACX framework in achieving this pivotal and informative feature learning outcome.

Using this approach, we refined the effects of drugs on the previously identified acceleration protrusion phenotype^6^ and discovered a novel protrusion phenotype termed bursting protrusion, specifically potentiated by the myosin inhibitor, blebbistatin. Myosin is known to influence leadin -edge dynamics through processes such as focal adhesion assembly, cortical contractility enhancement, and actin retrograde flow amplification, and previous reports indicate blebbistatin’s promotion of protrusions in both 2D^65^ and 3D^63,64^ environments. Our analysis indicates that pinpointing such specific subcellular phenotypes enhances our understanding of cell motility and morphodynamics, facilitating the characterization of precise drug effects at subcellular and cellular levels. Moreover, by scrutinizing subcellular phenotypes characterized by DeepHACX, we identified nine previously unknown single-cell dynamic phenotypes of protrusion dynamics, each exhibiting differential drug sensitivities. This underscores the importance of multi-scale analysis spanning subcellular to cellular scales, offering a more comprehensive understanding of the heterogeneity of drug actions. Our findings suggest an unexpectedly rich phenotypic diversity in cellular dynamics at subcellular and single-cell levels, emphasizing the potential of DeepHACX in unraveling intricate cellular behaviors.

In conclusion, DeepHACX represents a significant advancement in the field of live cell phenotyping, offering a unique combination of interpretability, heterogeneity-preserving feature learning, and concise representation. Its implications for drug discovery, cell biology, and personalized medicine make it a valuable tool for researchers seeking a deeper understanding of cellular dynamics in response to molecular perturbations.

## Methods and Methods

### Data Preparation

The cell culture and live cell imaging procedures were followed according to the previous studies (Lee, 2015, Wang, 2018). For the drug treatment experiments, we cultured PtK1 cells on 27mm glass-bottom dishes (Thermo Scientific cat. #150682) for two days and stained them with 55µgml^-1^ CellMask Deep Red (Invitrogen) following the manufacturer’s protocol. We monitored the cell using a spinning-disk confocal microscope. We visually examined cellular morphology, the level of protein expression, and the number of nuclei in each cell movie. We performed our analysis using the cells with a flat, minimally ruffling morphology and wide leading edges, the low expression level of fluorescent proteins, and a single nucleus. At this stage, we did not know the cluster distribution along the cell edges. Thus, this data selection can be assumed unbiased for the presented analyses.

For Arp2/3 inhibition experiments, cells were incubated with 50 µM of CK666 or CK689 (EMD Millipore) for an hour before imaging. For Cytochalasin D experiments, cells were incubated with DMSO or Cytochalasin D (50 or 100 nM) (Sigma) for half an hour before imaging. For myosin inhibition experiments, cells were incubated with 20 µM blebbistatin (EMD millipore, cat. # 023389) for a half-hour before imaging. In order to prepare a large dataset to train a deep learning model, we also included the data from the previous study in addition to the data generated in this study^6^, which gave us more than 24000 protrusion time-series.

### Local sampling and event registration

After the threshold-based segmentation, the cell edge velocity was calculated by tracking the cell edges^5,35,40^. Then, probing windows with the size of 500 nm by 500 nm were placed along the cell boundary^5^. The number of probing windows then maintained constant throughout the movie (∼100 windows in each cell). The local protrusion velocity and fluorescence intensity were quantified by averaging the values within probing windows. By repeating this procedure in each frame of the time-lapse movies, we acquired the time series of protrusion velocities and fluorescence intensities. Then, the time of protrusion onset was determined in each probing window^5^. The protrusion velocity and fluorescence intensities over time in individual windows were registered by aligning the protrusion onset at t = 0.

### Treating missing values

In the case where velocity values are missed from a time series due to image noise, these missing values are treated as follows: For each edge velocity or fluorescence intensity sample, the entire time series was discarded if the length of continuous missing values is longer than a threshold (here, we used the value 8). Otherwise, the average of four values before and after the missing value was used to individually estimate the value for this location.

### De-noising the samples by Empirical Mode Decomposition

For each registered sample, the edge displacement was calculated from the edge velocity using the Matlab function *trapz()*. Empirical Mode Decomposition (EMD)^69^ was then applied to the transformed protrusion edge displacement to remove noise. Finally, the denoised velocity was calculated from the denoised displacement using the Matlab function *diff()*.

Cell movement is highly non-stationary, and Empirical Mode Decomposition (EMD)^69^ is a local and data-driven de-noising method to decompose non-stationary signals into a series of intrinsic components. The general procedure of EMD can be described as follows:

1). Identify all the extremes (minima and maxima) of sample *d(t)*;
2). Connect the local maximum points and local minimum points separately using an interpolation method to generate the envelope, *e(t)*;
3). Compute the average of envelopes, *avg(t)* = (*e_min_*(*t*) + *e*_max_(*t*))/2;
4). Eliminate the average signal of the envelope from the sample *d(t)* to obtain the residual: *avg(t) = (e_min_*(*t*) + *e_max_*(*t*)/2;
5). Iterate from steps 1) to 4) on the residual *m(t)* until the *avg(t)* equals zero.

After EMD, the original signals can be decomposed into intrinsic mode functions (IMF) without any loss of information, and the residue is called the trend. For each component, a de-trended fluctuation analysis was used to measure the self-affinity as the fractal scaling index (α), which estimates the fractal-like autocorrelation properties. The value of α inversely correlated with the possibility that the component is originated from noise. In our procedure, the code was obtained from a previous publication^70^, and the value of α was empirically set to 0.33 to balance the maintaining information and trimming noise.

### Representing the velocity by Symbolic Aggregate approximation (SAX)

The dimensionality of times series samples consisting of 56 frames remained still high, and we were interested in general patterns over large time scales. Therefore, the dimensionality of time series data should be reduced by some dimension reduction methods. To this end, we applied SAX (Symbolic Aggregate approXimation)^71^ to our time series dataset to reduce dimensionality and discretize the data. The general procedure of SAX is summarized as follows:

1). Manually determine the reduced dimension, *N*, and the symbolic number, *M* (the number of discretization levels).
2). For each normalized sample, the time series data over the entire time range are pooled together and fitted to a Gaussian distribution. The entire time series is divided into *M* ranges with equal probabilities of the fitted Gaussian distribution. We represent the values of each range a symbol accordingly.
3). Subsequently, the time series is divided into *N* intervals along over time. The average value is calculated in each interval to represent the raw time series data.
4). Finally, the average value in each interval is represented by the symbol defined in step 2.
5). Iterate from step 2) and 4) until all samples are represented.

After the SAX representation, all time series data are reduced to low-dimensional (N) symbolic series data, which are used in further analyses. In the current experiment, M and N were both empirically set to four. Here, 4 symbols that range from 0 to 3 will be used to calculate the autocorrelation coefficients. In addition, as an implicit benefit, the symbol representation process in SAX also removes local noise.

### Calculating the sample dissimilarity

To measure the dissimilarity of two time series, the original description of SAX representation proposed an approximate Euclidean distance of SAX as a dissimilarity measure^72^. Instead, we used the dissimilarity measure based on estimated autocorrelation functions (ACF) ^73^. First, the estimated autocorrelation vector was calculated, and the square Euclidean distance between the autocorrelation coefficients was then used to measure the dissimilarity of two samples as follows:

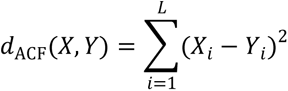

In our implementation, the ACF distance was calculated using the TSdist R package^74^.

### Unsupervised Teacher Model Using ACF feature

1. Calculation of partial similarity matrix

1. After the velocity time series are denoised by Empirical Mode Decomposition (EMD), we pooled the time series whose temporal lengths are similar within the threshold (6 frames; 30 seconds).
2. We pad random noise to the end of each time series to make the length equal to the longest one. We generate the random noise from the gaussian distribution with the mean estimated by the average value of the last five time points. Then, the missing part will be padded with random noise with the estimated mean and standard deviation.
3. To reduce the dimensionality of time series, we represented the time series data by Symbolic ApproXimate representation (SAX)^75^ as described in our previous study^6^.
4. We calculate the Euclidean distance based on the autocorrelation coefficient.
5. Repeat from steps 1) to 4) until all the samples are calculated.
6. The final distance similarity matrix is the average mean of distance similarity matrix and its transpose to guarantee that the similarity matrix is symmetric.

2. **Clustering:** We applied community detection method^55^ to the similarity matrix. First, we made a K-nearest neighbor graph based on similarity distance. Then, we calculated the adjacency matrix and identified the communities using the R package *igraph*. The number of clusters was estimated based on the silhouette values of clustering results.

### Student Model for Deep Feature Learning

**1. Velocity time series preprocessing.** In this step, we will perform nonlinear scaling of protrusion velocity to reduce the effect of large magnitude of protrusion velocity since the large magnitude may come from less accurate measurement. The majority of velocity magnitude should be less than 10µm/min based on our experience. Therefore, we manually designed a sigmoidal mapping function, 2/(1 + *e*^−0.3*v*^) – 1. After this sigmoidal scaling, the range of the velocity becomes [-1, 1].

**2. Model Training.** We randomly split the dataset into three parts: training, validation, and test sets with a ratio: 0.49, 0.21, 0.3. The training set was used to fit the parameters of the model, while the validation set was used to select the model with the best fit (the lowest value of the objective function). The details of our proposed supervised Bi-LSTM autoencoder were shown in Fig. 2a. Mainly we utilized three layers of bidirectional long short-term memory (bi-LSTM) as an encoder to extract the features and combined another three layers of bi-LSTM as a decoder to reconstruct the input. In order to make the representative features consistent with the clustering results from the ACF-based clustering, we added a multilayer perception (MLP) classifier to guide the training process. The total loss includes two loss functions as follows.

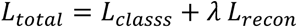

where reconstruction loss (*L_recon_*) is the mean squared error function from the autoencoder, the classification loss (*L_classs_*) is the multiple-categorical cross-entropy function from the MLP, *λ*is the balancing weight between *L_classs_* and *L_recon_*.

**3. Model Selection.** In order to optimize the balancing weight between cross-entropy for classifier training and mean squared error for autoencoder training, we trained the model with varying weights from 1 to 50 and selected the best weight that provides the most discriminative features for CK666 perturbation by maximizing effect size. We used a training set to fit the parameters with the batch size 128 and 237 epochs. During the training, we monitored the loss in the validation set and use the model parameters for the best performance with the validation set. We used the bi-LSTM encoder to extract the features for the subsequent analysis. TensorFlow was used to implement the supervised bi-LSTM autoencoder in Python 3 in Ubuntu 18.04.

### Phenotyping Using Deep Features

We extracted the features from the trained bi-LSTM encoder and then applied Principal Component Analysis (PCA) for the dimensional reduction of the learned features. Based on the percentage of the explained variance, the first 15 principal components are used for clustering analysis.

After the feature reduction, the dataset was split into paired experiments: CK689/CK666, DMSO/CyD50/CyD100, DMSO/Bleb. For each paired experiment, we calculated the sample similarity using Euclidean distance and then apply community detection on the selected samples with 51 frames. To evaluate the optimal number of clusters, we calculated silhouette values to estimate the optimal numbers in each experiment on the pooled control samples.

### Deep phenotyping Using Deep Features

After the initial clustering, we further sub-divided the phenotypes of bursting protrusion (Cluster II) and accelerating protrusion (cluster VI) into deep phenotypes using the deep features. We first pooled all the samples from the target phenotypes from different paired experiments and then applied the community detection to determine the deep phenotypes based on the Euclidean distance. The optimal number of clusters was determined by the maximum silhouette value.

### Interpretation of deep features

To elucidate contributions of protrusion velocities to deep phenotypes, we conducted feature differential expression analysis followed by SAGE^53^, a model-agnostic method designed to quantify predictive power while considering feature interactions. Initially, we identified deep features extracted from the BiLSTM encoder that were uniquely associated with each specific deep phenotype of Accelerating or Bursting phenotypes. This selection process utilized the Wilcox rank sum test, implemented through the presto R package. Subsequently, for the identified deep features, we employed the SAGE analysis^53^ to pinpoint time intervals significantly contributing to the prediction of the associated deep phenotypes. This was achieved using the framework of additive importance measures, employing Shapley values. The SAGE functions were implemented using the SAGE python package.

### Statistical Testing of Heterogenous Drug Effects

We quantified the drug effects on each phenotype based on its proportion. We counted the number of each phenotype in each cell for control and drug-treatment conditions. These numbers in each cell were resampled using *bootstrp()* in MATLAB to build 10,000 different bootstrapped datasets, and the distribution of the proportional differences of each phenotype between control and drug-treatment conditions were created. Using these distributions, p-values were calculated by estimating the probability that the phenotype proportion of one condition was greater or less than that of the other condition. The 95% confidence intervals of the proportions were estimated by the MATLAB *bootci()* function. This procedure does not assume the distribution of the data.

### Cluster Visualization

For each paired experiment, we applied t-SNE (t-distribution stochastic neighboring embedding) for visualization with the default parameter (PC number:15, perplexity:30). The sample densities on two-dimensional t-SNE mappings were estimated using the crowdedness of each sample below the radius threshold, which was implemented as scatplot in MATLAB by Alex Sanchez.

### Cellular Phenotyping based on subcellular phenotypes

We used the proportions of six identified subcellular phenotypes as the cellular morphodynamic features and then applied the previously described clustering method to identify cellular phenotypes from 133 cell movies. To quantify the drug effect on the cell phenotypes, we pooled Cell Cluster 6, 8, 9 into Accelerating Cell Group, Cell Cluster 3, 6, 7, 8 into Bursting Cell Group, and Cell Cluster 1, 2 into Steady Cell Group. Then, we calculated the proportion differences between Accelerating/Bursting and Steady Cell Groups. Also, we estimated the uncertainty of proportion difference using a random resampling strategy with replacement for 1000 times using the boot.ci function with the parameter “norm” in *boot* R package.

## Data availability statement

The datasets used in the current study are available from the corresponding author on a reasonable request.

## Code availability statement

The datasets used in the current study are available from the corresponding author on a reasonable request.

## Acknowledgments

We thank Microsoft for providing us with Azure cloud computing resources (Microsoft Azure Research Award), and Boston Scientific for providing us with the gift for deep learning research. This work was supported by NIH grants, R35GM133725, R01HL163513, and R15GM122012.

## Author Contributions

C.W. initiated the project, designed the algorithm of deep learning, performed the drug response analysis, and interpretation analysis, and wrote the final version of the manuscript and supplement. H.J.C. designed and performed the live cell imaging experiments, drug perturbation experiments; L. W. performed the clustering analysis of the time series of unequal temporal lengths; K.L. coordinated the study and wrote the final version of the manuscript and supplement. All authors discussed the results of the study

## Competing Interests

The authors declare no competing financial or non-financial interests.

## Author Information

Correspondence and requests for materials, data, and code should be addressed to K.L.

## Notes

### Competing Interest Statement

The authors have declared no competing interest.

### Summary of Updates

New Figure 6; Other figures and manuscript were extensively revised; Supplemental files updated.

