## Supplementary Figure, Supplementary Table for "Uncovering Interpretable Fine-Grained Phenotypes of Subcellular Dynamics through Unsupervised Self-Training of Deep Neural Networks"

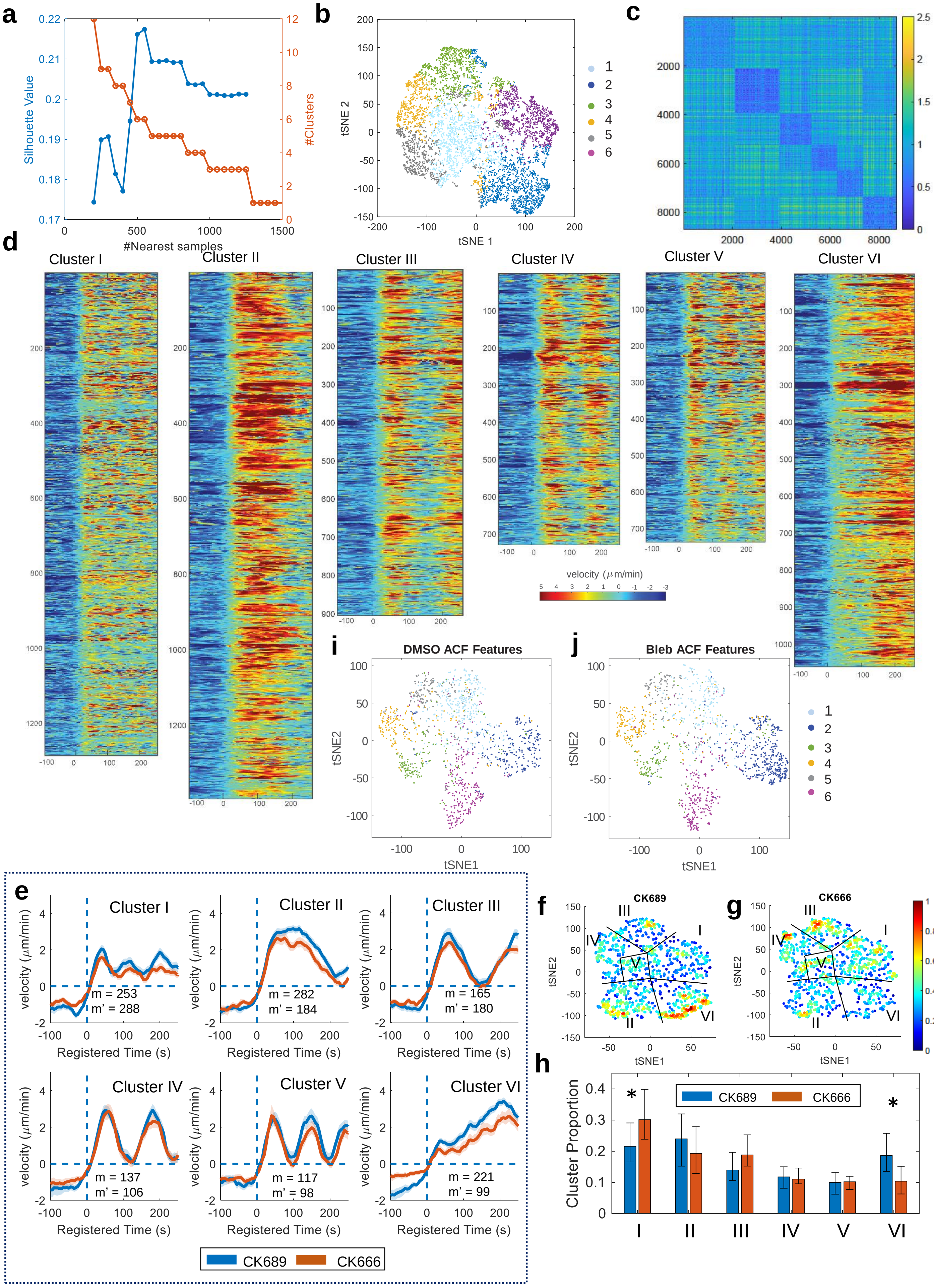

a
Silhouette Value
#Clusters
#Nearest samples
b
tSNE 1
tSNE 2
1
2
3
4
5
6
c
d
Cluster I
Cluster II
Cluster III
Cluster IV
Cluster V
Cluster VI
velocity (μm/min)
i
DMSO ACF Features
j
Bleb ACF Features
tSNE1
tSNE2
e
Cluster I
m = 253
m' = 288
Cluster II
m = 282
m' = 184
Cluster III
m = 165
m' = 180
Cluster IV
m = 137
m' = 106
Cluster V
m = 117
m' = 98
Cluster VI
m = 221
m' = 99
velocity (μm/min)
Registered Time (s)
CK689
CK666
f
CK689
g
CK666
h
Cluster Proportion
CK689
CK666
I
II
III
IV
V
VI

**Supplementary Figure 1. Subcellular protrusion phenotypes identified by the unsupervised teach model**. (**a**) The average Silhouette value and the number of clusters with the varying number of neighbors in the community detection clustering. (**b**) The t-SNE plot of the autocorrelation functions of protrusion velocity time series overlaid with cluster assignments. (**c**) The distance similarity heatmap ordered by the cluster labels. (**d**) The full protrusion velocity heatmaps in six identified phenotypes. (**e**) Averaged velocity time series in each cluster in CK689(control) and CK666-treated cells (m: the number of probing windows in CK689-treated cells; m': the number of probing windows in CK666-treated cells). (**f-g**) The t-SNE plot of ACFs overlaid with the data density and cluster assignments CK689 (f), CK666 (g). (**h**) Effects of CK666 on each protrusion phenotype. * $p<0.05$ indicates the statistical significance by bootstrap sampling. The numbers of cells: 10 for CK689 and 10 for CK666. (**i-j**) the t-SNE plot of ACFs overlaid with cluster assignments in DMSO (**i**) and blebbistatin-treated cells (**j**).

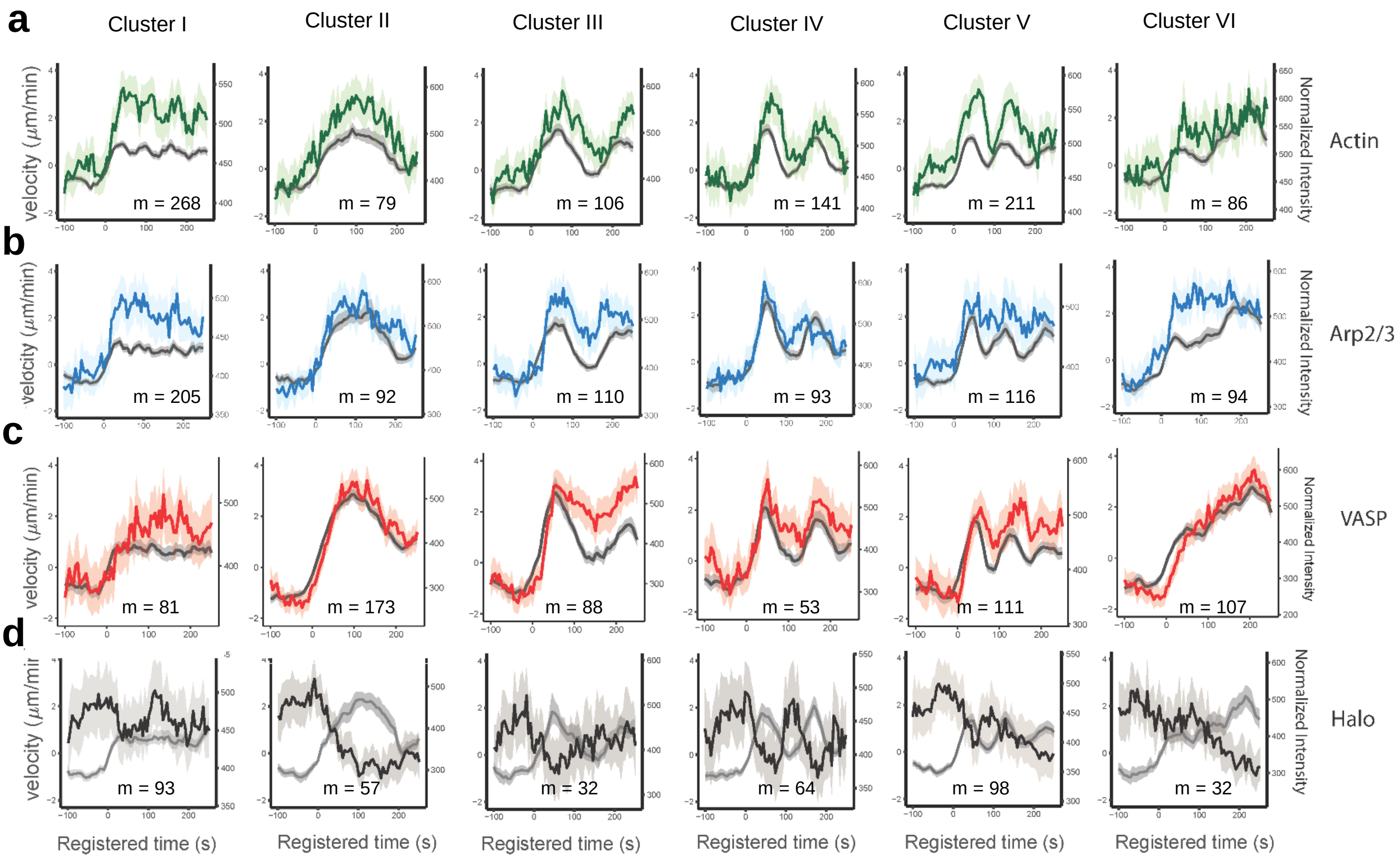


**Supplementary Figure 2. Differential actin regulator dynamics associated with subcellular protrusion phenotypes.** (**a–d**) Averaged normalized fluorescence intensity time series in each phenotype, overlapped with the corresponding velocity profiles. Time series are registered with respect to protrusion onset (t = 0). Solid lines indicate population averages. Shaded error bands indicate 95% confidence intervals of the mean computed by bootstrap sampling. The gray lines indicate protrusion velocity time series associated with the indicated fluorescent proteins. M: the number of probing windows. The numbers of cells: 10 for actin, 11 for Arp2/3, 9 for VASP, and 5 for Halo)

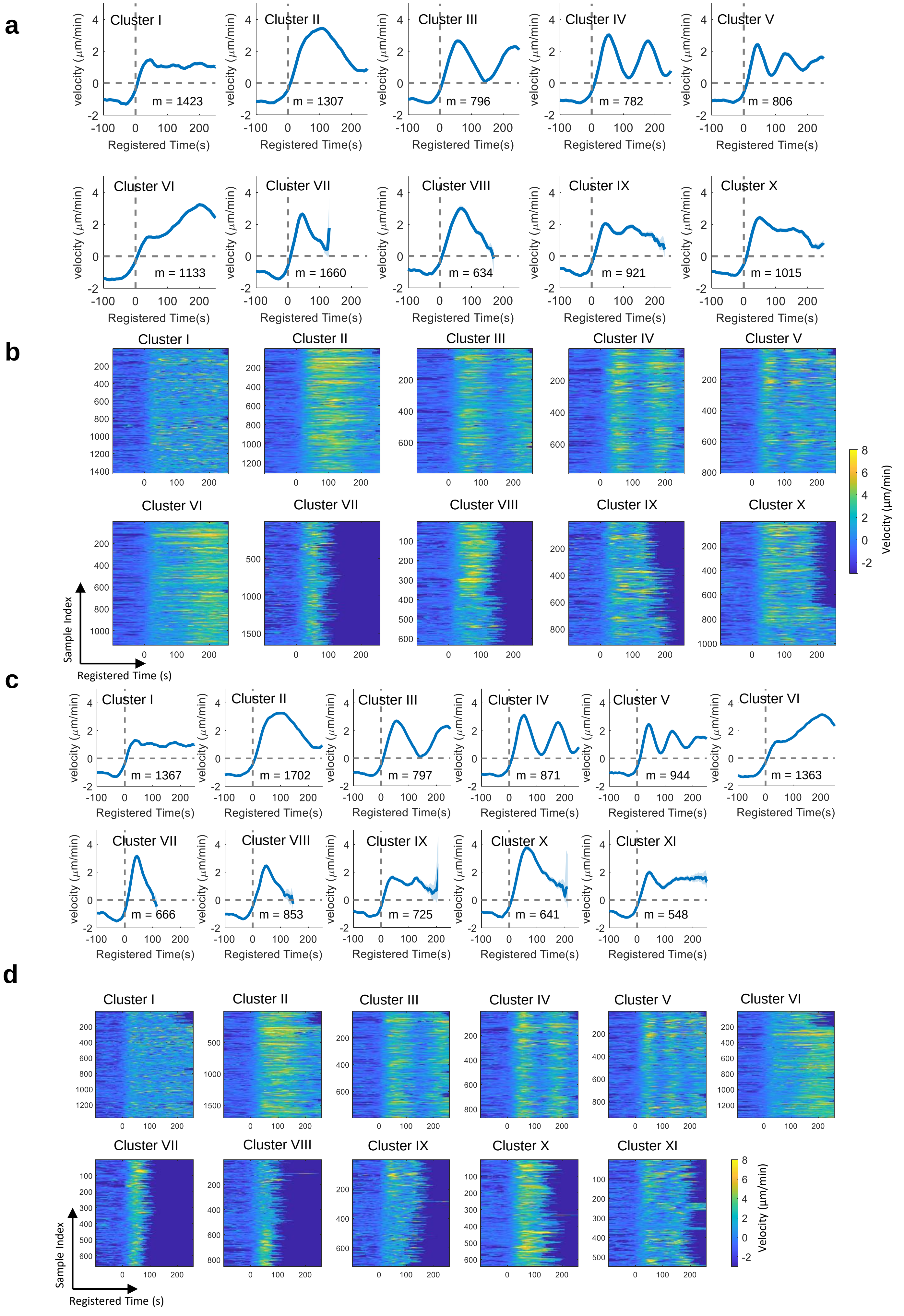
a
Cluster I
m = 1423
Cluster II
m = 1307
Cluster III
m = 796
Cluster IV
m = 782
Cluster V
m = 806
Cluster VI
m = 1133
Cluster VII
m = 1660
Cluster VIII
m = 634
Cluster IX
m = 921
Cluster X
m = 1015
velocity (μm/min)
Registered Time(s)
b
Cluster I
Cluster II
Cluster III
Cluster IV
Cluster V
Cluster VI
Cluster VII
Cluster VIII
Cluster IX
Cluster X
Velocity (μm/min)
Sample Index
Registered Time (s)
c
Cluster I
m = 1367
Cluster II
m = 1702
Cluster III
m = 797
Cluster IV
m = 871
Cluster V
m = 944
Cluster VI
m = 1363
Cluster VII
m = 666
Cluster VIII
m = 853
Cluster IX
m = 725
Cluster X
m = 641
Cluster XI
m = 548
velocity (μm/min)
Registered Time(s)
d
Cluster I
Cluster II
Cluster III
Cluster IV
Cluster V
Cluster VI
Cluster VII
Cluster VIII
Cluster IX
Cluster X
Cluster XI
Velocity (μm/min)
Sample Index
Registered Time (s)

**Supplementary Figure 3. Subcellular protrusion phenotypes from variable-length time series of protrusion velocities.** (**a-b**) Averaged velocity time series profiles in each protrusion phenotype based on ACF features (**a**) and the corresponding velocity heatmaps (**b**). (**c-d**) Averaged velocity time series profiles in each protrusion phenotype based on deep features extracted by the student deep model (**c**) and the corresponding velocity heatmaps (**d**). Solid lines indicate population averages and shaded error bands indicate 95% confidence intervals of the mean estimated by bootstrap sampling. The error bars indicate 95% confidence interval of the mean of the cluster proportions. m: the number of probing windows.

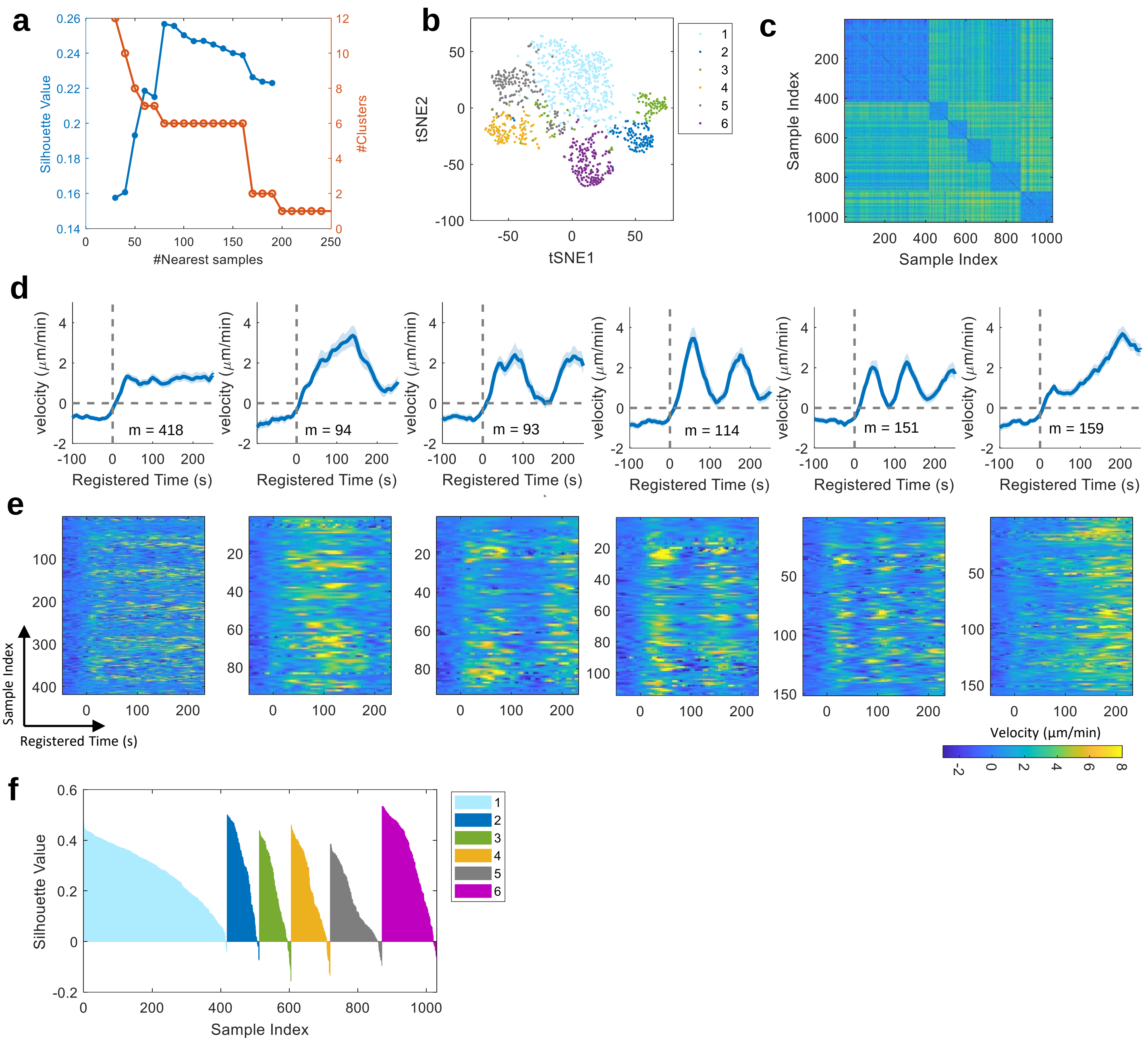


**Supplementary Figure 4. DF-based Subcellular protrusion phenotypes of MCF10A cells. (a)** The average Silhouette value and the number of clusters with the varying number of neighbors in the community detection clustering. (**b**) The t-SNE plot of the DFs of protrusion velocity time series overlaid with cluster assignments. (**c**) The distance similarity heatmap ordered by the cluster labels. (**d-e**) Averaged velocity time series profiles in each protrusion phenotype based on DFs extracted by the student model (**d**) and the corresponding velocity heatmaps (**e**). Solid lines indicate population averages and shaded error bands indicate 95% confidence intervals of the mean estimated by bootstrap sampling. **(f)** Silhouette value per each sample across different clusters. The *m* represents the number of probing windows.

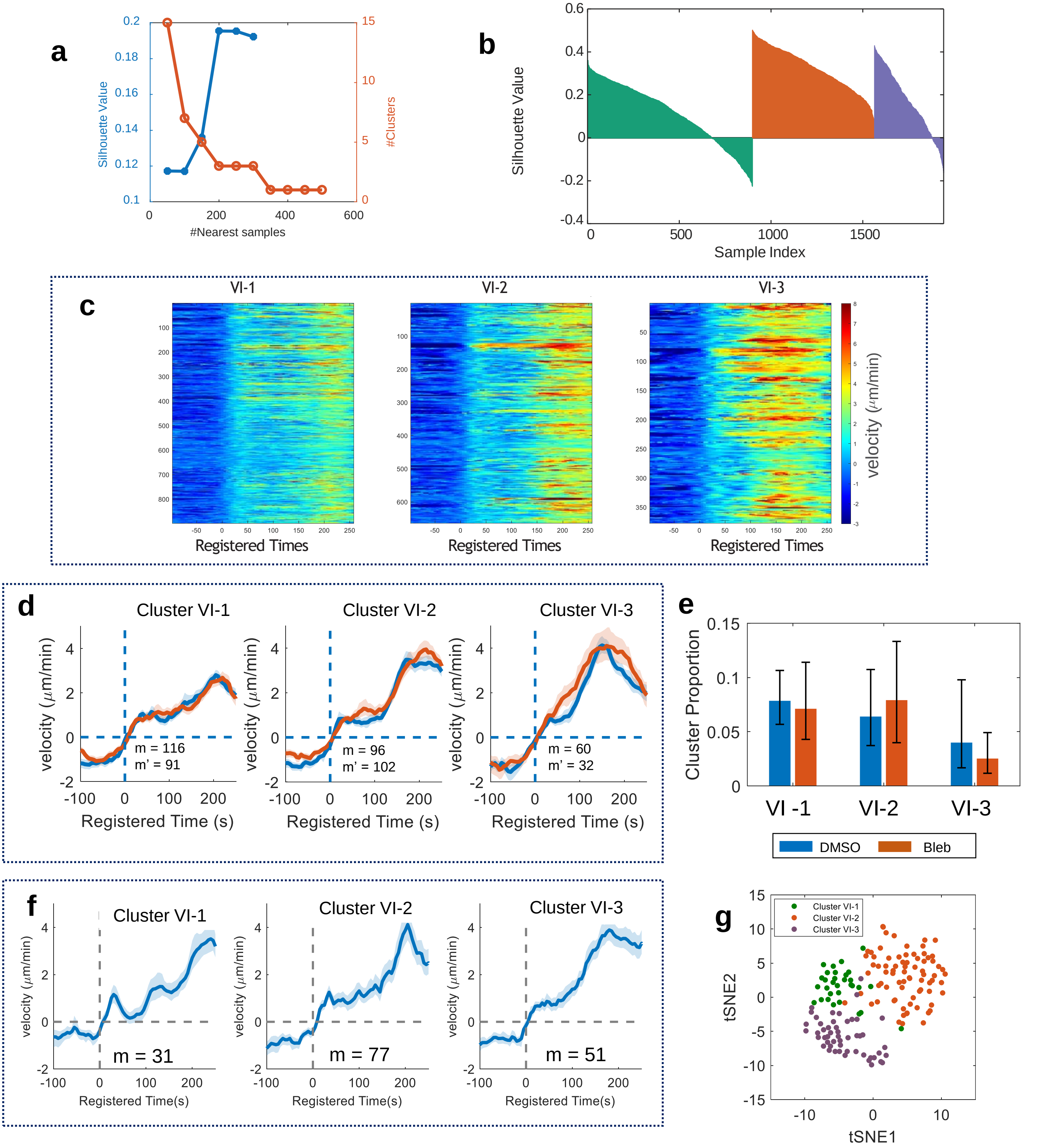


**Supplementary Figure 5. Sub-clustering results from the accelerating protrusion phenotype.** (**a**) The average Silhouette value and the number of clusters with the varying number of neighbors in the community detection clustering. (**b**) The silhouette plot the clustering result. (**c**) The velocity heatmap of three deep accelerating protrusion phenotypes. (**d**) Averaged velocity time series profiles in each deep phenotype in DMSO and blebbistatin-treated cells (m: the number of probing windows in DMSO; m': the number of probing windows in blebbistatin-treated cells). (**e**) Effects of blebbistatin on each protrusion phenotype. There was no statistical significance in each phenotype. The numbers of cells: 14 for DMSO and blebbistatin for 13. The error bars indicate 95% confidence interval of the mean of the cluster proportions. (**f-g**) Sub-clustering results of the accelerating protrusion phenotype of MCF10A cells. (**f**) Averaged velocity time series profiles in each deep phenotype. m: the number of probing windows. (**g**) The t-SNE visualization of the deep features inaccelerating protrusion phenotype. Solid lines indicate population averages and shaded error bands indicate 95% confidence intervals of the mean estimated by bootstrap sampling.

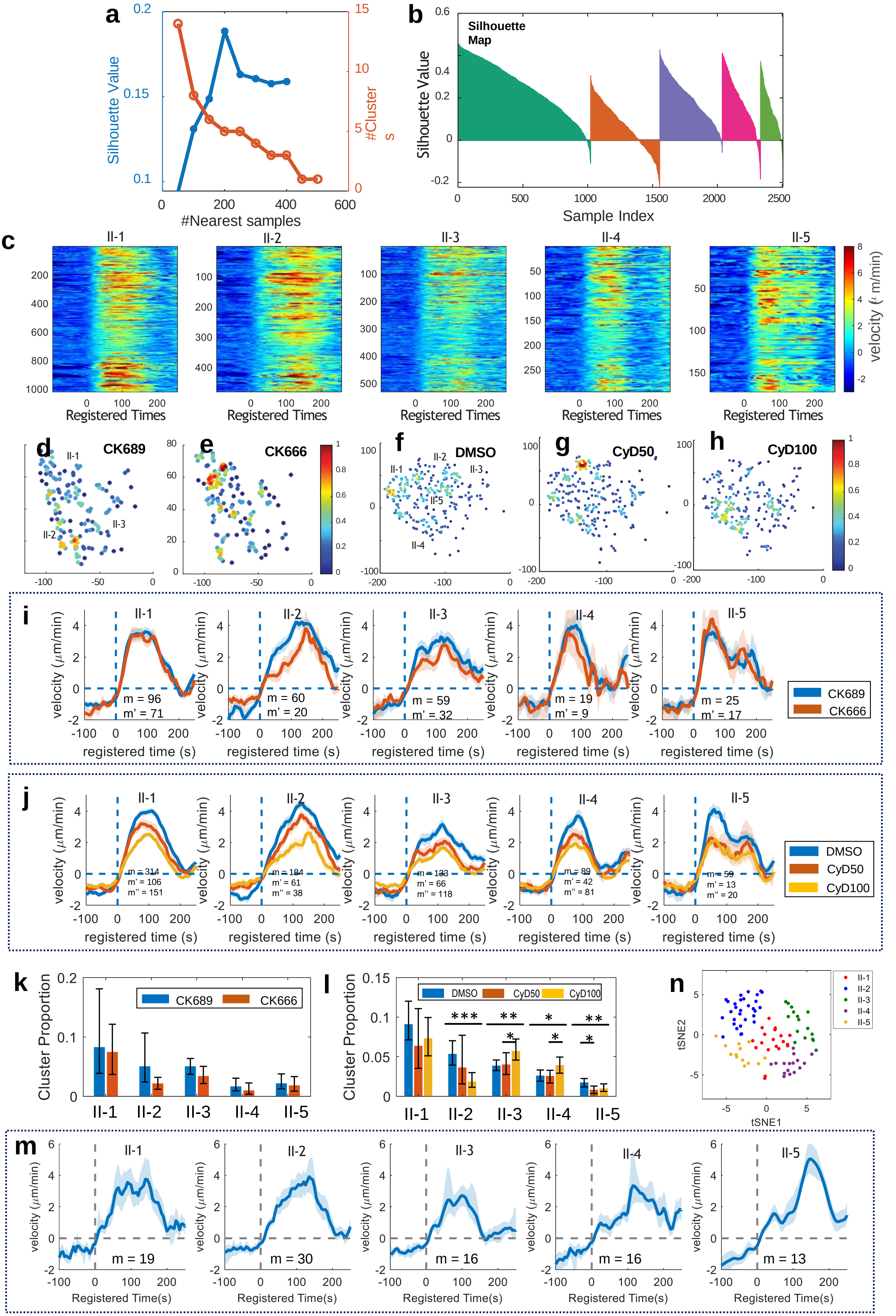

a
Silhouette Value
#Clusters
#Nearest samples
b
Silhouette Map
Silhouette Value
Sample Index
c
II-1
II-2
II-3
II-4
II-5
Registered Times
velocity (μm/min)
d
CK689
e
CK666
f
DMSO
g
CyD50
h
CyD100
i
velocity (μm/min)
registered time (s)
m = 96
m' = 71
m = 60
m' = 20
m = 59
m' = 32
m = 19
m' = 9
m = 25
m' = 17
CK689
CK666
j
m = 314
m' = 106
m'' = 151
m = 184
m' = 61
m'' = 38
m = 133
m' = 66
m'' = 118
m = 89
m' = 42
m'' = 81
m = 59
m' = 13
m'' = 20
DMSO
CyD50
CyD100
k
Cluster Proportion
l
m
Registered Time(s)
m = 19
m = 30
m = 16
m = 16
m = 13
n
tSNE1
tSNE2

**Supplementary Figure 6. Sub-clustering results from the Bursting protrusion phenotype.** (**a**) The average Silhouette value and the number of clusters with the varying number of neighbors in the community detection clustering. (**b**) The silhouette plot the clustering result. (**c**) The velocity heatmap of three deep bursting protrusion phenotypes. (**d-f**) The t-SNE visualization of the deep features of the bursting protrusion for CK666 (**d-e**) and Cytochalasin D (**f-h**) treated cells. The color indicates the density of data on the t-SNE plots. (**i-j**) Averaged velocity time series profiles in each deep bursting protrusion phenotype in CK689/CK666-treated cells (**i**) (m: the number of probing windows in CK689-treated cells; m': the number of probing windows in CK666-treated cells). and DMSO/Cytochalasin D-treated cells (**j**) (m: the number of probing windows in DMSO-treated cells; m': the number of probing windows in CyD50-treated cells; m'': the number of probing windows in CyD100-treated cells). (**k-l**) Effects of CK666 (**k**) and Cytochalasin D (**l**) on each protrusion phenotype. The error bars indicate 95% confidence interval of the mean of the phenotype proportions. *p <0.05, ** p < 0.01, *** p < 0.001 indicate the statistical significance by bootstrap sampling. The numbers of cells: 10 for CK689, 10 for CK666 (i-k) and 22 for DMSO, 16 for CyD50, 20 for CyD100 (j-l). (**m**-**n**) Sub-clustering results of the bursting protrusion phenotype of MCF10A cells. (**m**) Averaged velocity time series profiles in each deep phenotype. m: the number of probing windows. (**n**) The t-SNE visualization of the deep features in burstng protrusion phenotype. Solid lines indicate population averages and shaded error bands indicate 95% confidence intervals of the mean estimated by bootstrap sampling.

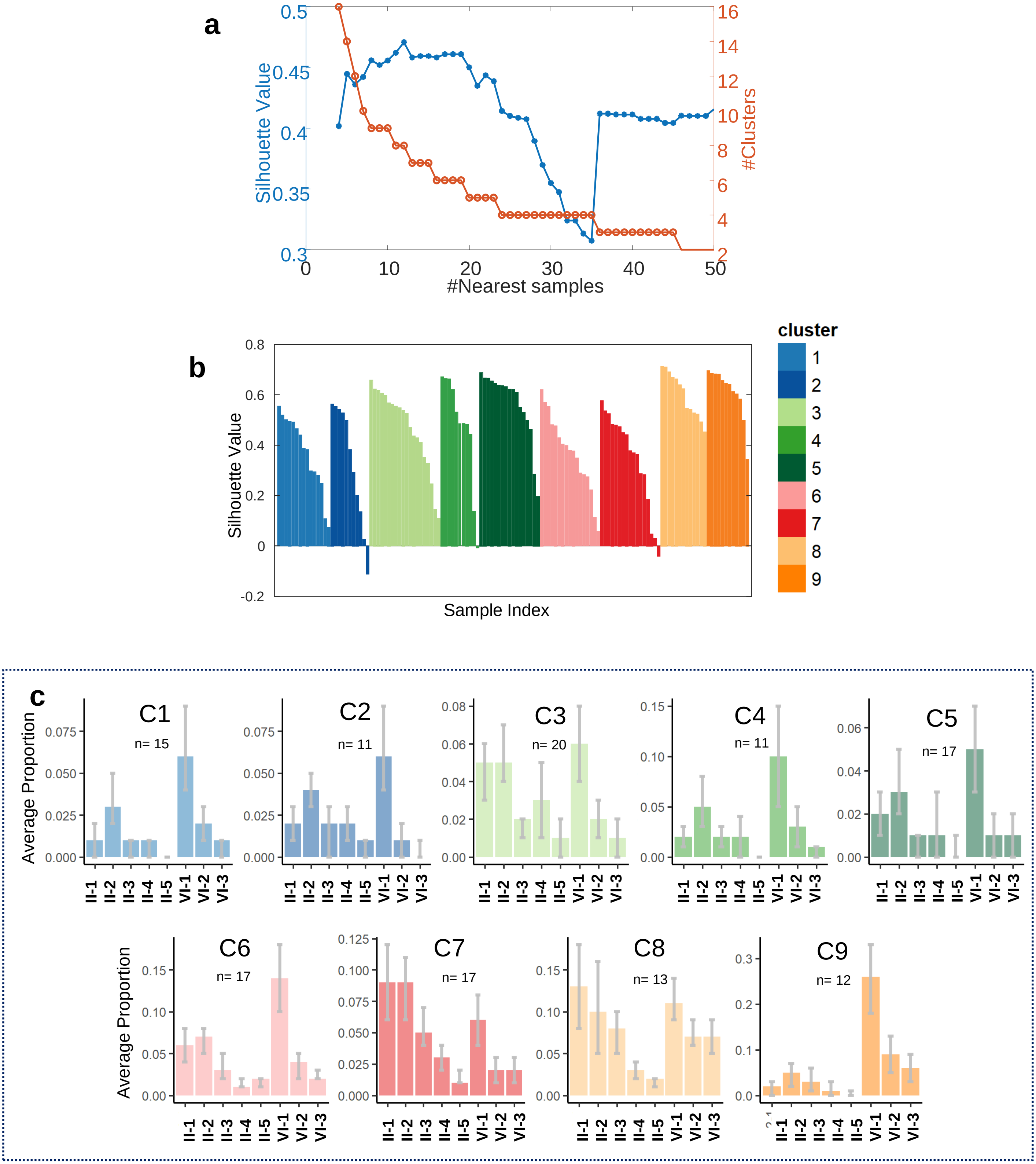


**Supplementary Figure 7. Cellular protrusion phenotyping.** (**a**) The average Silhouette value and the number of clusters with the varying number of neighbors in the community detection clustering. ((**b**) The silhouette plot the clustering result. (**c**) The proportions of deep phenotypes in each cell phenotypes (n: the number of cells in each cell cluster). The 95% confidence intervals of the mean were estimated by bootstrap sampling.

### Supplementary Table 1. Summary of single-cell protrusion phenotypes of PtK1 cells

| Cell Cluster # | Dominant Protrusion | Cell Phenotype | Drug-Sensitive Deep Phenotypes | |
|---|---|---|---|---|
| | | | Cluster II-1 (blebbistatin) | Cluster VI-2/3 (CK666/CyD) |
| 1 | Steady | Steady Cell1 | Low | Low |
| 2 | Steady | Steady Cell2 | Low | Low |
| 3 | Steady & Periodic #1 | Periodic Cell1 | **Mid** | Low |
| 4 | Steady & Periodic #2 | Periodic Cell2 | Low | Low |
| 5 | Steady & Periodic #3 | Periodic Cell3 | Low | Low |
| 6 | Steady, Bursting, & Accelerating | Mid-Bursting/ Accelerating Cell | **Mid** | **Mid** |
| 7 | Bursting | Bursting Cell | **High** | Low |
| 8 | Bursting & Accelerating | Strong-Bursting/ Accelerating Cell | **High** | **High** |
| 9 | Accelerating | Accelerating Cell | Low | **High** |
